# Disease-associated genetic variants can cause mutations in tissue-specific protein isoforms

**DOI:** 10.1101/2025.07.24.666385

**Authors:** Giovanna Weykopf, Mihaly Badonyi, Elias T Friman, Jasmine Minh Hang Nguyen, Alexis Ioannou, Benjamin J Livesey, Audrey Coutts, Elizabeth F Hird, Murray Wham, Chloe M Stanton, Veronique Vitart, Jing Su, Lee Murphy, J. Kenneth Baillie, Mark D Gorrell, Joseph A Marsh, Wendy A Bickmore, Simon C Biddie

## Abstract

Genetic variants can cause protein-coding mutations that result in disease. Variants are typically interpreted using the reference transcript for a gene. However, most human multi-exon genes encode alternative isoforms. Here, we show that coding exons in alternative isoforms harbour more population variants than exons of reference isoforms, consistent with their reduced evolutionary constraint, and that these variants are more likely to cause nonsynonymous coding mutations. Common and rare disease-associated variants mapping to alternative transcripts can lead to amino acid substitutions predicted to be structurally damaging in the corresponding protein isoform. The alternative transcripts to which disease-associated variants map demonstrate high tissue-specific expression, with many unannotated in reference human genomes, revealed only by long-read RNA-sequencing. As an example, we report an unannotated alternative transcript of the inflammasome regulator DPP9 that is lung epithelium-specific and which harbours a common genetic variant associated with severe COVID-19 and lung fibrosis. The variant causes a p.Leu8Pro missense mutation in an alternative first exon, predicted to disrupt the encoded alpha helix. These findings highlight the importance of considering alternative isoforms, their tissue-specific expression, and full-length transcripts in variant interpretation, with implications for uncovering underappreciated mechanisms of both common and rare disease.

## Main

Transcript isoforms are alternative mRNAs produced from a single gene locus, through mechanisms such as alternative splicing or the use of alternative promoters (Kjer-Hansen et al., 2024). Approximately 95% of human multi-exon genes generate multiple transcript isoforms, thereby expanding protein diversity (Pan et al., 2008). Protein-coding alternative isoforms show high tissue-specific expression (Glinos et al., 2022) and can be developmental stage-specific or context-dependent (Robinson et al., 2021). Alternative isoforms can have different functions than the main isoform, including gain-of-function or dominant-negative properties (Yang et al., 2016). For example, transcription factor isoforms can have differential binding activities (Lambourne et al., 2025), while immune receptor isoforms can have dominant-negative effects on the immune response (Pasquesi et al., 2024).

Rare genetic variants that alter isoform expression or sequence are known to contribute to disease. Splice site mutations, for example, can lead to developmental and neuropsychiatric disorders by disrupting isoform balance or intron inclusion (Barbaux et al., 1997, Fernando et al., 2025). However, most clinical interpretation of coding variants remains limited to the reference isoform (Richards et al., 2015), and only isolated cases of pathogenic variants specific to alternative isoforms have been reported. One example is a missense mutation in an adult-specific isoform of *SCN5A*, associated with cardiovascular conductive disease (Veerman et al., 2017). Recent advances in long-read RNA sequencing have revealed previously unannotated transcripts and highlighted the extent of tissue-specific isoform expression (Glinos et al., 2022, Reese et al., 2023). Thus, there is a need to re-examine disease-associated mutations in this broad isoform landscape.

Although many alternative transcripts are detected, the majority may not encode stable or functional proteins. For most highly expressed protein-coding genes, one isoform predominates expression (Ezkurdia et al., 2015), and many alternative isoforms lack cross-species conservation or signs of purifying selection (Tress et al., 2017, Pozo et al., 2021). Analyses of human genetic variation similarly suggest that most alternative exons evolve under relaxed constraint (Liu and Lin. 2015). These observations imply that, while some alternative isoforms may be biochemically competent, they are often not required for function and may represent non-essential or non-adaptive transcriptional byproducts. A key unmet need is to determine which genetic variants exert their effects through specific alternative isoforms, particularly those expressed in restricted tissues or developmental stages.

Here, we address this challenge by integrating large protein language model-based variant effect prediction (VEP), AlphaFold structural modelling and tissue-specific long-read transcriptomic data to prioritise alternative isoforms expressed in relevant biological contexts. We focus on common and rare genetic variants that occur in protein-coding exons specific to alternative isoforms (‘alt-exons’), often in a tissue-specific manner, identifying missense mutations that are predicted to be conserved or structurally damaging. As a case study, we analyse fine-mapped GWAS variants associated with severe COVID-19 (Pairo-Castineira et al., 2023) and identify a common variant located in an alternative first exon of an unannotated, lung cell-specific isoform of *DPP9,* causing a leucine to proline substitution within an isoform-specific N-terminal region, predicted to disrupt an alpha-helix. Taken together, our observations reveal a largely unexplored mechanism by which common and rare disease-associated variants can act through coding changes in tissue-specific alternative isoforms, frequently absent from reference annotations.

## Mutations in alternative isoform-specific exons

To determine the contribution of genetic variants to coding mutations in alternative isoforms, we first classified exons by their relation to reference transcript isoforms. We considered transcripts from a meta-analysis of long-read RNA-sequencing (RNA-seq) based on GTEx, ENCODE and other datasets (Shi et al., 2024), extracted exons, and classified these into exons present in reference transcript isoforms (ref-exons), or exons specific to alternative transcript isoforms (alt-exons). Reference transcript isoforms were defined using the Matched Annotation from NCBI and EMBL-EBI (MANE) Select transcript (Morales et al., 2022). Although exons in the MANE Select transcripts can also appear in alternative isoforms, we classify all exons present in the MANE Select transcripts as ref-exons (**figure 1A**). Alt-exons were further classed into those annotated in Ensembl (exons in catalogue; EIC), or unannotated exons from novel long-read RNA-seq transcripts (exons not in catalogue; ENIC). Across the human genome, there were over 200k ref-exons, while alt-exon counts were approximately 170k for EIC and 180k for ENIC (**figure 1B**). We classified EIC relative to the reference transcript into: alternative first exons (AFEs), alternative internal exons, alternative last exons, alternative 5’ splice sites, alternative 3’ splice sites or single exons (**figure 1A**). For EIC, the most common were 5’ and 3’ splice site extensions, followed by AFEs (**figure S1A**). Considering all exons across all transcripts from a gene, we observe a medium number of seven exons per gene (**figure S1B**).

**Figure 1.**
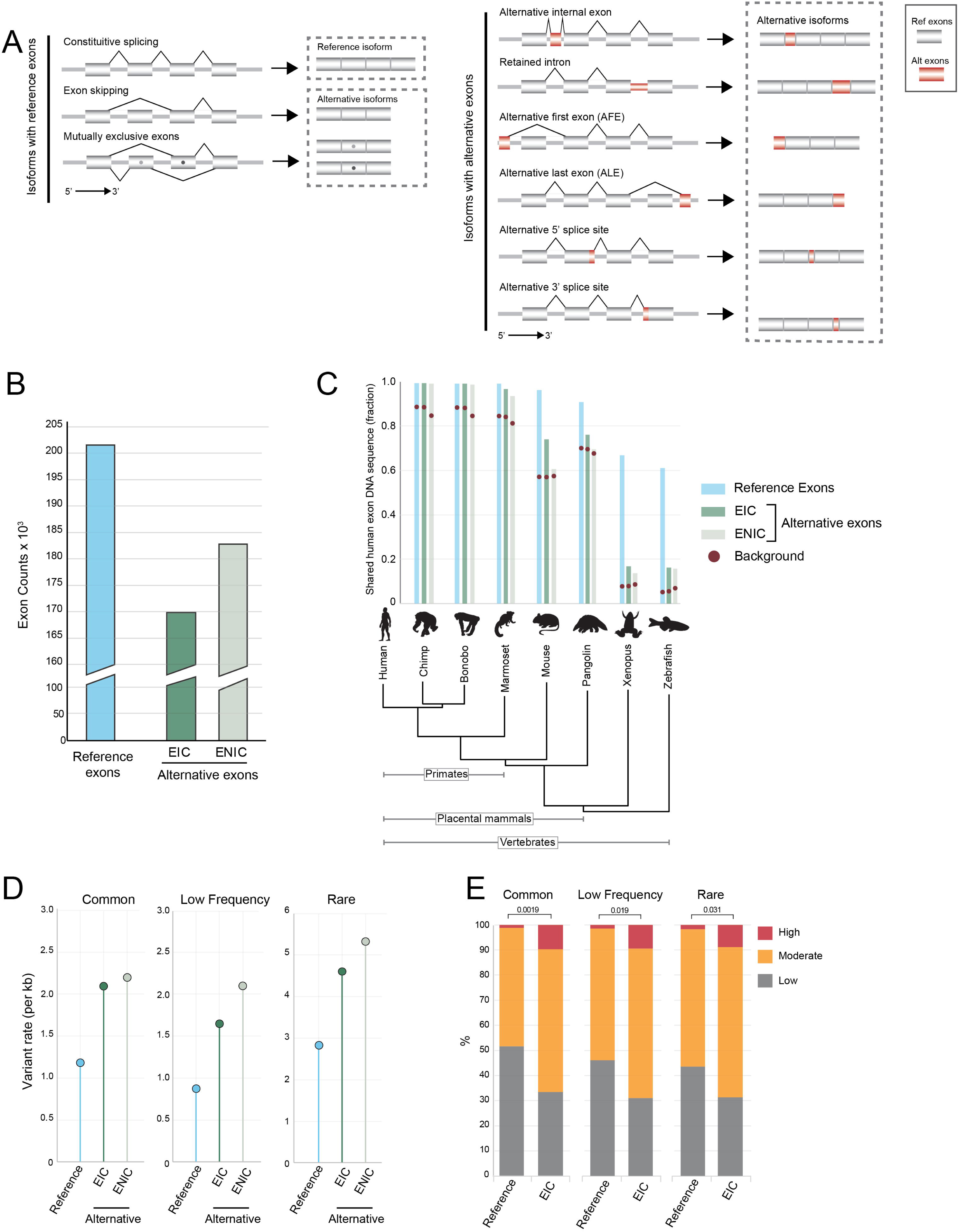
Features and classification of human exons. **A.** Schematic of isoform structures, depicting ref-exons consisting of exons present in the reference isoform, and mechanisms generating alternative isoforms from ref-exons. Isoforms containing alt-exons (red), which are exons specific to alternative isoforms, can be formed through multiple mechanisms. **B.** Bar graph of exon counts for ref-exons, and alt-exons, divided into exons in catalogue (EIC) and exons not in catalogue (ENIC) for human assembly Hg38. **C.** Evolutionary conservation of exons (ref-exon, EIC and ENIC) was determined by the fraction of exons per class shared between other species, including primates, placental mammals and evolutionarily distant vertebrates. Exons that were considered shared required a minimum match of 50% of nucleotides using LiftOver. Background models were generated using shuffled nucleotide sequences for the exons in each class, with the mean of 5 imputations depicted. **D.** Variant rate (per kilobase; kb) for each exon class was determined using single nucleotide variants (SNV) from GnomAD (v4). Variants were grouped by minor allele frequency (MAF): common variants, >0.05 to < 0.5; low frequency variants, > 0.01 <0.05; and rare >0.001 and < 0.01. The number of variants for each exon class were normalised by the combined exon genomic length to determine the per kb rate. **E.** Percentage bar chart of SNV impact on coding sequences using isoform-aware annotation for Ensembl transcripts. Variants were mapped to ref-exon and alt-exon EIC, grouped by allele frequency from gnomAD (v4). Predicted coding variant annotations were grouped into low (synonymous), moderate (missense), and high (frameshift, stop gain, stop lost and start gain). Where variants are annotated to multiple transcript isoforms, the most severe coding impact was considered. P-values were calculated using ANOVA of synonymous versus nonsynonymous proportions.

To consider evolutionary conservation as a mark of functional exons (Márquez et al., 2021, Keren et al., 2010), we performed a pairwise analysis of exon classes using sequence orthology with LiftOver, requiring a minimum base fraction of 0.5 (**figure 1C**). Ref-exons and alt-exons were observed to have a high fraction of orthologous exons in primates, but alt-exons were less conserved than ref-exons in placental and more distant vertebrate comparisons. Ref-exons have positive mean PhyloP scores, while alt-exons have near neutral mean PhyloP scores (**figure S1C).** Some alt-exons may have evolved more recently, being primate- or human-specific. Transposable elements, a class of repetitive sequences, have been proposed to contribute to accelerated genomic regions (Feschotte, 2008).

Consistent with this, ref-exons have a lower fraction of repetitive sequence-harbouring exons compared to alt-exons (**figure S1D**), with Short Interspersed Nuclear elements (SINEs), and Long Interspersed Nuclear Elements (LINEs) representing the highest proportion in alt-exons. Within the SINE family, the primate-specific Alu elements, which have been implicated in exonization events (Krull et al., 2005), were most abundant (**figure 1SE**). Together, these data show that alt-exons have recently evolved and many may have originated from retroelement transposition.

To understand the contribution of genetic variants to protein-coding regions, we analysed the population prevalence of genetic variants in exons, using aggregated allelic frequencies from gnomAD v4.1.0 (Chen et al., 2024). We grouped single nucleotide variants (SNVs) by their minor allele frequency (MAF) into common, low frequency, and rare, observing comparable distribution of variants in exon classes (**figure 1SF**). Across variant frequency groups, variant rate (variants per kilobase) for alt-exons was higher (1.7 – 2.4-fold) than for ref-exons (**figure 1D**). We next considered the impact of variants on protein-coding sequences, using isoform-aware variant annotation in Ensembl annotated isoforms. We grouped the variant impacts into low (synonymous), moderate (missense), and high (including frameshift, stop gain, stop loss and start gain), and took the highest severity for a variant where the variant occurs in multiple transcript isoforms (**figure 1E**). For common variants in ref-exons, the majority (51.6%) were synonymous, while most common variants in alt-exons were missense mutations (56.8%). Across all MAF groups, common to rare, the proportion of variants causing moderate and high impact were significantly higher in alt-exons than ref-exons. Thus, alt-exons have a higher variant burden and higher proportion of damaging variants.

## Disease-associated variants in tissue-specific alternative isoforms

Genetic variants associated with disease, whether from GWAS or rare variant databases, are rarely interpreted in the context of transcript isoforms (Cummings et al., 2020), even though many alternative isoforms show tissue-specific expression and can encode distinct proteins. Therefore, we investigated the extent to which disease-associated variants map to alt-exons and whether these events may have functional consequences.

From the GWAS Catalog (Cerezo et al., 2025), we find ∼40,000 variants mapping to alt-exons across ∼15,000 alternative transcript isoforms from GTEx long-read RNA-seq data from 22 GTEx tissues. Using Gini coefficients to quantify tissue specificity (Kryuchkova-Mostacci and Robinson-Rechav. 2017), we found that most alternative isoforms showed more tissue-specific expression than their matched reference isoform (**figure 2A**). While alternative isoforms are typically expressed at lower levels, 32% exceeded the expression of the reference in at least one tissue (**figure 2B**), consistent with previous estimates (Yang et al., 2016). Grouping GWAS traits into biological systems, for example clustering metabolic or immunological-associated traits, we find that GWAS variants are enriched in alt-exons across biological systems (**figure S2A**). Additionally, variant-associated alternative transcript isoforms were highly expressed in relevant tissues. For example, variants associated with lung, endocrine, or haematological traits map to alternative isoforms that are expressed in diseases-relevant tissues (**figure 2B, S2B**).

**Figure 2.**
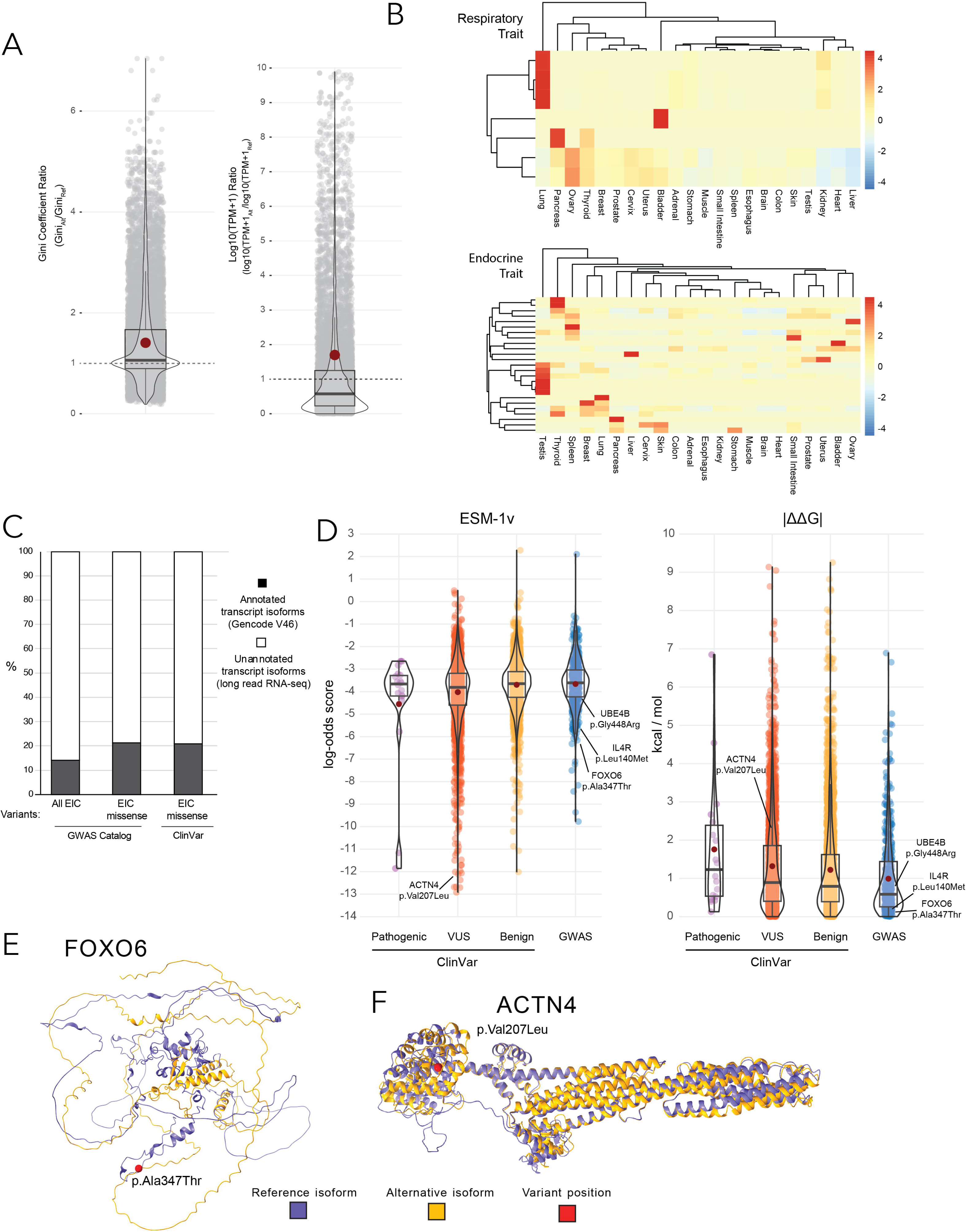
Common and rare variants are associated with tissue-specific isoforms. **A.** Box and violin plots of alternative isoform: reference isoform ratios for tissue-specificity metric (Gini coefficient ratios, left), and transformed expression (log10(transcripts per million (TPM)+1) ratios, right). GWAS variant-mapped alternative isoforms and gene-matched reference isoforms were annotated with Gini coefficients and TPM values from long-read RNA-seq of 22 GTEx tissues. For transformed TPM, ratios were determined from the highest expression in any GTEx tissue. Mean is shown in red. **B.** Heatmap of alternative transcript isoform expression for isoforms associated with grouped GWAS variant traits. Expression is from long-read RNA-seq transcripts in 22 GTEx tissues. Expression as z-score of log10 TPM. Hierarchical clustering for tissues and variants was performed using the ward method. **C.** Proportional bar chart of the number of Gencode (v46) annotated and unannotated isoforms for transcripts associated with variants from GWAS catalog and ClinVar in non-canonical exons in catalogue (EIC). **D.** Violin and box plots of evolutionary scale modelling (ESM)-1v and FoldX calculated protein thermodynamic stability difference in the Gibbs free energy (ΔΔG) scores. Predicted ClinVar and GWAS catalog missense variant impact in variant-associated alternative isoforms were based on predicted Alphafold 3 (AF3) structures. ClinVar variants were grouped by ClinVar classifications into pathogenic, variants of uncertain significance (VUS) and benign. ClinVar variants were grouped by ClinVar annotated pathogeniticity into pathogenic, variants of uncertain significance (VUS) and benign. **E** and **F.** Reference isoform (blue) and variant-associated alternative isoform (yellow) for (E) FOXO6 with GWAS variant position shown in red, or for (F) ACTN4 with ClinVar variant position shown in red.

Next, we sought to investigate the impacts of disease-associated variants in alternative isoforms, considering the GWAS Catalog variants, as well as pathogenic, uncertain significance and benign variants from ClinVar. To identify coding variants affecting alternative protein isoforms, variants were annotated against isoform-specific EIC coordinates and mapped to coding transcripts derived from long-read RNA-seq data (**figure S2C**). Of the alternative transcript isoforms that harbour variants mapping to an alternative isoform-specific exons, ∼80% were unannotated transcripts (**figure 2C**), reinforcing the importance of long-read transcript discovery. Similar to GWAS variants, alternative isoforms with mapped ClinVar variants showed high tissue-specificity, although most have lower expression than the matched reference isoform (**figure S2D**). We further considered only those variants that lead to missense changes in alternate exons, excluding those predicted to disrupt splicing and transcripts predicted to undergo nonsense-mediated decay (NMD).

To estimate the functional consequences of each missense variant, we used ESM-1v (Meier et al., 2021), a large protein language model that predicts the effect of amino acid substitutions based solely on sequence context, avoiding the necessity of deep sequence alignments, required by most variant effect predictors (VEP), that limit applicability to alternate isoforms. To consider protein structural impacts of missense variants, we used AlphaFold 3 (AF3) to model the structures of ∼17,000 alternative isoforms. We then assessed the structural impact of each variant by estimating changes in Gibbs free energy of folding (ΔΔG) with FoldX (Delgado et al., 2019). We consider the absolute ΔΔG due to the observation that this better reflects pathogenicity (Gerasimavicius et al., 2020), likely because both stabilising and destabilising variants can cause disease. As expected, ClinVar pathogenic variants are the most damaging according to both metrics (**figure 2D).** However, this group is small, with only 18 pathogenic variants mapping to missense changes in alternative isoforms, likely reflecting variants not annotated as coding in alternative isoforms often being overlooked in clinical interpretation. The next most damaging group comprises ClinVar variants of uncertain significance (VUS), consistent with the expectation that some will be clinically pathogenic. ClinVar benign and GWAS variants are predicted to have the mildest impacts. This may represent GWAS variants that are not causal due to linkage disequilibrium, or exert relatively modest effects compared to rare variants implicated in Mendelian disease.

We highlight several cases in which disease-associated variants map to tissue-specific alternative isoforms and are predicted to be damaging. A common GWAS variant associated with vitamin D levels (Hendi et al., 2023) maps to a disordered region of an unannotated *FOXO6* isoform (**figure 2E**) that is highly expressed in ovaries (**figure S2E**). A rare GWAS variant linked to eosinophil counts falls within an unannotated *IL4R* transcript (**figure S2F**) that is expressed in lung and spleen (**figure S2E**). A rare variant associated with atrial fibrillation (Roselli et al., 2018, Verma et al., 2024) affects a known UBE4B isoform (**figure S2G**), with expression in heart and muscle (**figure S2E**), consistent with previous observations (Mammen et al., 2011). Among ClinVar VUS, we identify a missense variant in an alpha-helical region of an unannotated ACTN4 isoform (**figure 2F**) expressed in brain, kidney and testes (**figure S2E**). *ACTN4* variants are associated with focal segmental glomerulosclerosis (Henderson et al., 2009), suggesting possible relevance.

We also find four VUS in an alternative exon of *EYA4* (Eyes Absent 4), with high sequence homology to a downstream ref-exon (**figure S3A**) and encoding a conserved protein tyrosine phosphatase domain (de la Peña Avalos et al., 2024) (**figure S3B**). These exons undergo mutually exclusive splicing, and the homology has likely arisen from a duplication event (Martinez-Gomez et al., 2022). The predicted reference and alternative isoforms have structural homology (**figure S3C**), and VEP scores for the VUS in the alternative isoform suggest impact damaging effects (**figure S3D**). Pathogenic ClinVar EYA4 variants are associated with cardiomyopathy and hearing loss, and the alternate isoform has tissue-restricted expression, detected in muscle, heart, and cervix (**figure S3E**). Details of these variants and the associated alternative isoforms coding mutations are detailed in supplemental **table S1**.

## A common disease-associated variant in an alternative DPP9 isoform

To exemplify coding mutations in alternative isoforms associated with a specific phenotype, we used GWAS fine-mapping of severe COVID-19 variants (Pairo-Castineira et al., 2023), consisting of 1,848 variants at a posterior probability of 95%. We find 178 variants in coding regions, 98 of which occur in alt-exon EICs, 4.1-fold above the expected genomic distribution (**figure S4A**). The EIC variants in alt-exon have a range of MAFs, most of which (94/98) have a GWAS p-value of > 1×10^−15^ (**figure 3A**). One common variant chr19:4717660-A>G (rs12610495, MAF=0.29, homozygous frequency 0.033), with a p-value of 9×10^−51^, is the lead variant at the *DPP9* locus (Pairo-Castineira et al., 2023) and has previously been tagged in GWAS for lung fibrosis (Fingerlin et al., 2013). *DPP9* encodes a serine protease that cleaves N-terminal dipeptides (Geiss-Friedlander et al., 2009, Ajami et al., 2004) and forms a complex with, and thereby suppresses, NLRP1 - the primary inflammasome and viral sensor in barrier epithelial cells (Barry et al., 2023, Nguyen et al., 2025, Zhong et al., 2018). Loss or inhibition of DPP9 function leads to inflammasome activation and cell death by pyroptosis (Okondo et al., 2017, Zhong et al., 2018). Rare coding variants in the reference isoform have been associated with inflammatory disease, including respiratory manifestations (Harapas et al., 2022, Wolf et al., 2023). Co-localization analysis suggests chr19:4717660-A>G is causative for COVID-19 and lung fibrosis (Dalapati et al., 2025) and has been previously suggested as an eQTL (Dalapati et al., 2025), and sQTL (Nakanishi et al., 2023) for *DPP9*.

**Figure 3.**
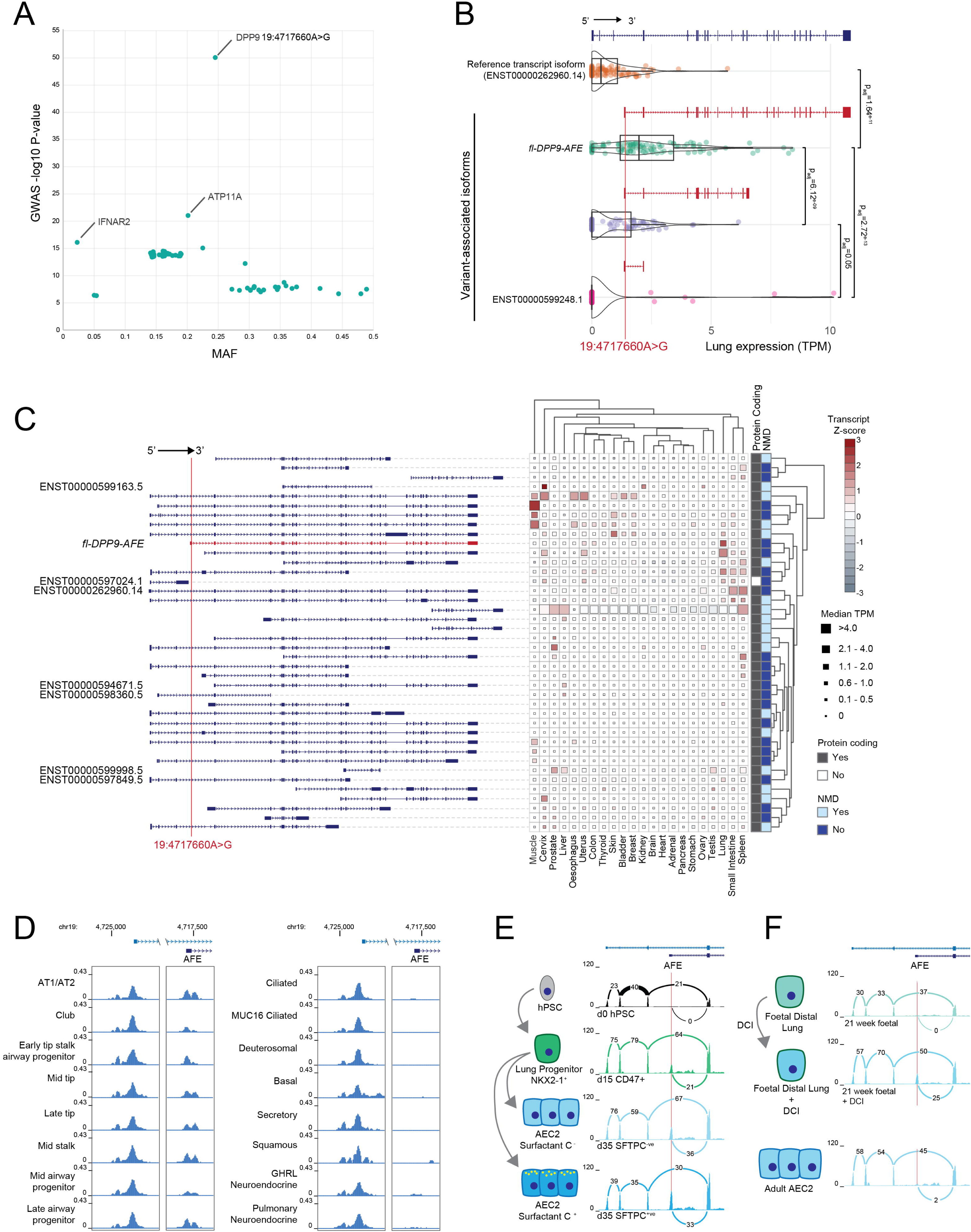
A variant-associated *DPP9* alternative isoform is expressed in lung. **A.** Scatter plot of lead variants and 95% credible variant set from a meta-analysis of severe COVID-19 GWAS that intersect alt-exons in catalogue (EIC), plotting minor allele frequency (MAF) from gnomAD (v4) against the GWAS p-value (-log10). The top three variants by GWAS p-value are annotated. **B.** Violin and box plot for *DPP9* lung transcript expression (median transcript per million reads (TPM)) from long-read RNA-seq (GTEx) for the reference transcript, and transcript isoforms intersecting 19:4717660A>G (rs12610495): full length *DPP9* alternative first exon (*fl-DPP9-AFE*) isoform, an unannotated *DPP9-AFE* isoform, and the annotated *DPP9-AFE* transcript (ENST00000599248.1 in Gencode V46). **C.** Heatmap of *DPP9* transcript isoforms expression for 22 tissues from GTEx long-read RNA-seq. Only transcripts with median expression > 0 for any tissue are shown. Z-scores are depicted per transcript, calculated across tissues. TPM, predicted protein coding and nonsense-mediated decay (NMD) are shown. Hierarchical clustering of transcript and tissues used the ward method. **D.** Genome browser image of published single-cell ATAC-seq tracked for epithelial cell types from foetal human lung (He et al., 2022). Pseudo-bulk ATAC-seq signal for the promoter of the reference transcript (left) and alternative promoter of the 19:4717660A>G variant-associated transcripts isoform (right). **E.** Expression of *DPP9* splice junction reads of human pluripotent stem cells (hPSC) from published short-read RNA-seq, differentiated into NKX2-1 positive lung progenitor cells, and surfactant C-positive and negative type-2 alveolar epithelial cells (AEC2). Tracks indicate normalised mapped reads from short-read RNA-seq, with numbers indicating normalised exon-exon spanning read counts. **F.** Expression of *DPP9* splice junction reads of foetal distal lung and adult AEC2 cells from published short-read RNA-seq. Foetal distal lung cells were induced to differentiation using dexamethasone, cyclic AMP, and 3-isobutyl-1-methylxanthine (DCI).

While the chr19:4717660-A>G variant has been frequently annotated as intronic to *DPP9*, we find it in an alt-exon. It intersects one GENCODE (v46) *DPP9* transcript (ENST00000599248.1), annotated as a short two exon non-coding transcript, where the alt-exon forms an AFE. Using *DPP9* transcript isoforms from long-read RNA-seq (Shi et al., 2024), we find three transcript isoforms that share the variant-harbouring AFE. The lack of previous annotation of these transcripts is likely attributable to short-read sequencing being unable to resolve transcripts due to the shared 3’ exons. In GTEx long-read RNA-seq of lung (n = 100), we observe high expression of one of the three AFE transcripts (**figure 3B**). A future GENCODE release will model the ENST00000599248 transcript with the coding sequence (CDS) extended to full length. Here, we refer to this full-length (fl) transcript as *fl-DPP9-AFE*. Compared to the reference *DPP9* transcript, the *fl-DPP9-AFE* transcript has higher expression in lung (**figure 3B**). It shares 19 exons with the reference transcript that encodes functional cytoplasmic DPP9 and only differs in the AFE, which forms an N-terminal extension. Using 22 tissues with GTEx long-read RNA-seq, we find 40 *DPP9* transcripts with detectable expression, with 33 unannotated in GENCODE. For these 40 transcripts, we observe high tissue-specific transcript isoform expression, with *fl-DPP9-AFE* being specific to lung (**figure 3C**). However, not all of these 40 transcripts are likely to encode proteins with 1/40 predicted to be non-coding, and 18/40 predicted to undergo NMD (**figure 3C**).

We also identified the *fl-DPP9-AFE* transcript isoform in published long-read RNA-seq datasets from a lung epithelial cell line, A549 (Chen et al., 2025). To independently validate *DPP9* transcript isoforms using deep targeted long-read RNA-seq, we developed Full-Length targEted capture using eXon probes for Isoform expRession (FLEXIR)-seq. This utilises a custom panel of commercial exon probes, optimized for Oxford Nanopore Technologies (ONT) long-read sequencing of polyA transcripts (**figure S4B**), covering all exons of the reference *DPP9* isoform, plus a probe for the *DPP9* AFE. Probes were additionally designed for neighbouring genes to increase library complexity. FLEXIR-seq enriched for target genes, with off-target genes representing fewer overall reads (**figure S4C**), and associated with high ranking and high expression in short-read total RNA-seq (**figure S4D**). FLEXIR-seq replicates were highly correlated (Pearson coefficients 0.88-0.99) demonstrating strong reproducibility (**figure S4E**). Relative expression of on-target genes showed good correlation with short-read total RNA-seq (R^2^=0.76), which we did not observe for off-target genes (R^2^=0.005) (**figure S4F**). Using this method, we confirmed the expression of *fl-DPP9-AFE* in lung epithelial cell lines A549, H358, HSAEC1-KT, HBEC3-KT and in N/TERT-1, a keratinocyte cell line (**figure S5A**). The *fl-DPP9-AFE* constituted 4 – 30% of total *DPP9* across cell lines (**figure S5A**). We further validated expression by qRT-PCR in A549, H358 and HSAEC1-KT cells, using exon-exon spanning primers that target the *DPP9-AFE* transcript (**figure S5B**).

As the chr19:4717660-A>G-variant-associated *fl-DPP9-AFE* transcript has lung-specific expression, we hypothesized that the AFE would utilise an alternative promoter, as previously observe for AFE isoforms (Baek et al., 2007). Using ENCODE data from foetal lung, the promoter of the reference *DPP9* isoform shows accessible chromatin associated with a CpG island, and DNaseI footprints suggesting transcription factor occupancy (**figure S5C**). The 5’ regulatory element of *fl-DPP9-AFE* also showed open chromatin, and DNaseI footprints, suggestive of an alternative promoter (**figure S5C**). In published single-cell ATAC-seq data from human foetal lung tissue (He et al., 2022), we observe an ATAC peak at the *fl-DPP9-AFE* alternative promoter in a subset of lung epithelial cell types from pseudo-bulk data (**figure 3D**), not accessible in mesenchyme, endothelial or immune cell types (**figure S5D**). The promoter of the reference transcript shows an ATAC peak in all cell types. To determine when during lung epithelial development *fl-DPP9-AFE* becomes expressed, we interrogated published short-read RNA-seq (Jacob et al., 2017) from lung epithelial differentiation of induced human pluripotent stem cells (hPSC). Taking exon spanning reads of the reference isoform and *fl-DPP9-AFE*, we detect no *fl-DPP9-AFE* expression in hPSC (**figure 3E**). Upon differentiation into lung progenitor cells and lung epithelial cells, *fl-DPP9-AFE* transcript expression emerged and increased during this trajectory, suggesting expression early in lung epithelial differentiation. There was minimal expression of *fl-DPP9-AFE* in quiescent foetal distal lung cells and adult alveolar epithelial type 2 (AT2) cells, but this increased in response to epithelial differentiation using dexamethasone, cyclic AMP, and 3-isobutyl-1-methyxanthine (DCI) in distal lung cells (**figure 3F**). These observations demonstrate that in lung, the *fl-DPP9-AFE* isoform is lung epithelial specific, with increased expression during lung epithelial differentiation.

## A DPP9 isoform-specific variant causes a missense mutation

The reference isoform of DPP9 has two translation initiation sites (Ajami et al., 2004), the first generating an 892 amino acid (aa) *DPP9-long* isoform preferentially located in the nucleus through an N-terminal nuclear localisation signal (Justa-Schuch et al., 2014), and the second generating an 863 aa *DPP9-short* isoform localised to the cytosol (Abbott and Gorrell. 2000, Ajami et al., 2004). DPP9-short specifically has been shown to inhibit NLRP1 inflammasome activation (Hollingsworth et al., 2021). *fl-DPP9-AFE* shares all exons encoding the cytoplasmic DPP9-short isoform, but with a predicted 33 aa N-terminal extension translated from the chr19:4717660-A>G variant-harbouring AFE. The AFE is in-frame with downstream exons and has an ATG start codon. Using a proteomic meta-analysis database (Fenyö and Beavis. 2015), we detect expression of the *fl-DPP9-AFE* encoded N-terminal peptide, enriched in lung (**figure 4A, S6A**), consistent with observations from *fl-DPP9-AFE* transcript expression.

**Figure 4.**
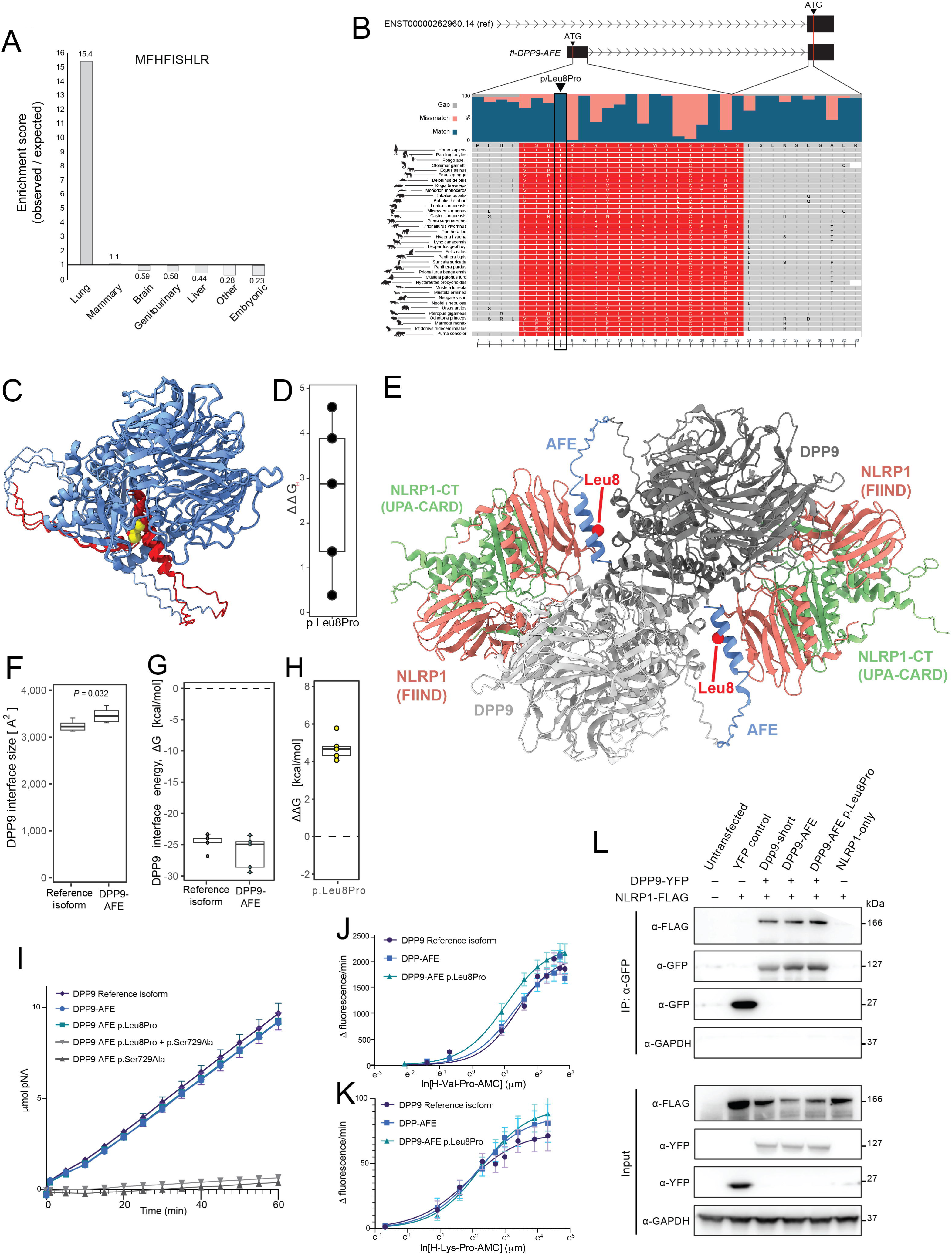
DPP9 common variant causes an isoform-specific N-terminus missense mutation. **A.** Enrichment of the N-terminal peptide sequence encoded by the DPP9-AFE of the 19:4717660A>G-associated transcript isoform in proteomic meta-analysis. Enrichment score was calculated by the distribution of peptide-expressing tissues, over a background tissue distribution, a composite of the five highest expressed reference isoform peptides (figure S3A). **B.** Multiple sequence alignment for amino acid conservation of the DPP9 alternate isoform AFE-encoding peptide using BLASTp. **C.** AlphaFold 3 (AF3) predicted structure of the DPP9 alternative isoform. DPP9 reference isoform in blue, N-terminus from the alternative isoform in red and p.Leu8Pro missense variant location in yellow. **D.** Variant effect prediction of the p.Leu8Pro missense mutations using AF3 predicted structure and FoldX calculated protein thermodynamic stability measured as difference in the Gibbs free energy (ΔΔG). Boxplot showing the FoldX-predicted ΔΔG of p.Leu8Pro across the AF3 models. **E.** AF3 predicted ternary structure of the DPP9 alternative isoform-NLRP1 complex, consisting of a DPP9 dimer, and two NLRP1 molecules. DPP9 reference isoform in grey, DPP9 N-terminus of the alternative isoform in blue, NLRP1 FIIND (Function-to-Find domain) in orange, and NLRP1 C-terminus (CT), termed UPA-CARD (UNC5, PIDD, and Ankyrin-caspase recruitment domain), in green. **F.** AF3-predicted structure interface size between the N-terminus of the DPP9 alternative isoform and NLRP1 using FreeSASA. **G.** AF3-predicted structure interaction energy between all chain pairs in the 2(NLRP1^full-length^– NLRP1^C-terminus^–DPP9) complex computed using FoldX 5.0 applied to the reference and alternative isoforms. **H.** FoldX estimations of ΔΔG from p.Leu8Pro missense mutations introduced into the N-terminus of the alternative DPP9 isoform of the AF3-predicted 2(NLRP1^full-length^–NLRP1^C-terminus^–DPP9) complex. **I.** DPP9-specific enzyme activity time course of DPP9 isoforms, with the p.Leu8Pro variant and, as controls, catalytically dead (p.Ser729Ala) variant, in whole cell lysates using the H-Gly-Pro-pNA dipeptide substrate at saturating concentration (1 mM). **J.** and K. DPP9-specific enzyme kinetics for DPP9 reference and AFE isoforms, and the p.Leu8Pro variant form of DPP9-AFE, using H-Val-Pro-AMC (J) and H-Lys-Pro-AMC (K) dipeptide substrates. Fluorescence emission was detected upon cleavage of the fluorogenic salt 7-amino-4-methylcoumarin (AMC), with rate of hydrolysis fit to an allosteric sigmoidal model. **L.** Western blot of co-immunoprecipitation for FLAG-tagged NLRP1 and yellow fluorescent protein (YPF)-tagged DPP9 reference and alternative isoforms, including the p.Leu8Pro mutation in the alternative isoform. Constructs were expressed in HEK293 cells, then immunoprecipitated with GFP antibodies to pull down YFP-tagged NLRP1.

The N-terminal DPP9 extension appears to have emerged within the Boreoeutheria magnaorder (**figure S6B,C**). In comparison, the DPP9 catalytic domain is highly conserved (Ross et al., 2018) across the mammalian lineage, with orthologs also detected in the fungal kingdom (**figure S6D**). The common variant, within the N-terminal extension, is predicted to cause a p.Leu8Pro missense mutation at a site that is highly conserved since its emergence in mammalians (**figure 4B**).

Applying AF3 to the *fl-DPP9-AFE* amino acid sequence, the predicted structure resembles the recently solved cryo-electron microscopy structure of human DPP9-short (Hollingsworth et al., 2021), with the N-terminal extension forming an alpha-helix (**figure 4C**). We used FoldX on AF3 structures, which suggests a helix-breaking impact of the leucine to proline substitution (**figure 6D**). The solved ternary DPP9-NLRP1 complex consists of two DPP9, each with two NLRP1 molecules - full-length NLRP1 and a C-terminus NLRP1 fragment (Hollingsworth et al., 2021). Using AF3 to model the ternary complex, with two DPP9 alternative isoform molecules (**figure 4E**), we find the N-terminal alpha-helix likely forms additional contacts with NLRP1 (**figure 4F**), increasing complex stability (**figure 4G**) compared to the reference isoform. Introducing p.Leu8Pro into the DPP9-AFE ternary complex with NLRP1 predicts disruption of these structural features (**figure 4H**).

DPP9 functions as a serine protease through a C-terminal α/β hydrolase domain. Assaying the enzymatic activity of the DPP9-AFE isoform and mutation effects, we find that DPP9-AFE has similar enzymatic activity to the reference DPP9-short protein in an *in vitro* H-Gly-Pro peptide hydrolysis assay (Ajami et al., 2004), and this is unaffected by the p.Leu8Pro mutation at saturating substrate concentrations (**figure 4I**). DPP9 can hydrolyse multiple proline-containing dipeptides. To explore substrate affinities, enzymatic kinetics were measured using substrates with differing charge and hydrophobicity. We find p.Leu8Pro DPP9-AFE to have a modest higher affinity (lower K_half_) for H-Val-Pro (**figure 4J**), but similar affinities for other dipeptides (**figure 4K, S6E,F**). To validate the predicted structural interactions of the DPP9 alternative isoform with NLRP1, we performed co-immunoprecipitation of YFP-tagged DPP9 isoforms and FLAG-tagged NLRP1 in HEK293T cells. Both the wild-type and p.Leu8Pro mutant DPP9 alternative isoforms showed interactions with NLRP1, comparable to those observe with DPP9-short (**figure 4J**), supporting the structural predictions.

## Discussion

Identifying the underlying mechanisms for both common and rare genetic variants remains a significant challenge. Variants impacting protein function are typically considered in relation to the reference isoform (Lek et al., 2016). Here we show the prevalence of coding exons in alternative transcript isoforms that are not annotated in the reference human genome, and are often highly tissue specific. While rare disease-causing coding variants in Mendelian disease are typically functionally damaging (Gerasimavicius et al., 2022), we demonstrate that some common variants in isoform-specific exons can cause similar events, disrupting the structure of isoform-specific protein regions. This highlights an underappreciated mechanism in variant interpretation, where we observe a discernible mutational effect for common and rare variants specific to alternative isoforms. Our framework can identify functional variants, and prioritise functional alternative isoforms, distinguishing these from the broader background of transcript noise (Varabyou et al., 2021).

Our observation of alt-exons having a higher proportion of transposable elements, in particular Alu elements, is consistent with the evolution of novel human or primate-specific exons by transposable element mediated exonization (Möller-Krull et al., 2008). Transposable elements are also a known source of genetic variation (Arribas et al., 2024, Ng and Xue, 2006, Pasquesi et al., 2024, Sela et al., 2010). We observe a higher variant burden and greater proportion of missense mutations in alt-exons. The persistence of variants in alt-exons is likely through either recent evolutionary emergence, or from weaker purifying selection compared to ref-exons, due to tissue-specific expression of the variant-harbouring isoforms, or buffering by other functionally compensating isoforms.

Amongst variants associated with severe COVID-19, we identified one within an unannotated, functional and lung epithelial-specific DPP9 isoform. The GWAS missense variant in this isoform is predicted to be structurally damaging, disrupting a predicted N-terminal alpha helix. DPP9 is the only known endogenous inhibitor of NLRP1, the primary inflammasome in barrier epithelial cells (Barry et al., 2023). Upon detection of pathogen-associated danger signals, NLRP1 initiates an inflammatory response resulting in pro-inflammatory cytokine secretion and pyroptosis. SARS-CoV-2 infection has been shown to cause NLRP1 activation in lung epithelial cells (Planès et al., 2022), consistent with the lung tissue type in which we find the variant-harbouring DPP9 isoform. The DPP9-NLRP1 axis is an important regulator of lung inflammation. Patients with rare loss-of-function mutations in DPP9 present with various immune disorders mediated by NLRP1 overactivation (Harapas et al., 2022), while an asthma-associated variant in NLRP1 (p.Met1184Val) has been reported to enhance binding of NLRP1 to DPP9 (Moecking et al., 2021, Moecking et al., 2022). The chr19:1717660-A>G *DPP9* variant has been reproducibly associated with both severe COVID-19 and lung fibrosis across multiple GWAS studies (Allen et al., 2020, Fingerlin et al., 2013, Pairo-Castineira et al., 2021, Pairo-Castineira et al., 2023), suggesting a unified mechanism of pulmonary inflammation and epithelial dysfunction. Functional enzymatic assessment of the DPP9-AFE p.Leu8Pro mutation suggests altered affinity for some dipeptides, suggesting favourable entrance of hydrophobic substrates into the catalytic pocket.

Alternative isoforms can differ in molecular interactions and function (Kelemen et al., 2013, Lambourne et al., 2025, Yang et al., 2016). We observe tissue-specific expression of variant-harbouring alternative isoforms in disease-relevant tissues. We leveraged AF3-predicted structures to generate two VEP scores and highlight various examples of likely structurally-damaging variants impacting isoform-specific protein regions. Inclusion of tissue-specific exons into protein structures is known to modify protein interactions (Buljan et al., 2012), thus coding variants in alt-exons could confer tissue-specific effects.

Discrepancies between reference annotations and the diversity of alternative isoforms has been reported (Morillon and Gautheret. 2019) and has resulted in both false positive and negative clinical diagnosis of variants later re-annotated as coding variants in alternative isoforms (Schoch et al., 2020). Long-read sequencing of relevant cell types or tissues is required to avoid mis-annotation of transcripts, as demonstrated here for *DPP9.* We validated our findings by developing a full-length targeted capture RNA-seq method, FLEXIR-seq, to allow deep characterization and quantification of full-length polyA transcripts. The method is analogous to similar methods, but has been optimized for ONT long-read sequencing and exon probes, which have been reported to provide uniform coverage to preserve relative quantification (Yaldiz et al., 2023).

Our study has several limitations. First, we used consortia long-read RNA-seq atlases (Glinos et al., 2022) to determine tissue-specific expression and association with disease variation, but these datasets do not consider disease contexts. As disease-specific long-read transcriptomics emerge (Gandal et al., 2018, Humphrey et al., 2025), taking context-specific transcript isoform expression levels into account could improve identification of functional isoforms, and variant interpretation and mechanistic insight (Cummings et al., 2020).

Additionally, full-length transcriptomes have not been resolved at the single-cell level, thus our variant interpretation likely misses transcripts expressed in rare cell types. Second, for complex traits with common variant association, we used the GWAS catalog which has limitations in reported SNP-trait associations, either inherent to individual studies such as sample sizes or ancestry, or where reported summary statistics likely miss causal variants in linkage disequilibrium. In this case, fine-mapped variants provide greater precision, and our analysis of a credible set of severe COVID-19 suggests enrichment of variants in alt-exons. A similar approach for other complex traits may identify more variants that operate through coding mutations in alternative isoforms. Third, we focused on missense variants in our analysis of disease-associated variants and structure-based variant effect predictions. This represents the largest class of nonsynonymous coding mutations, and a challenging coding mutation class to interpret, compared to, for example, loss-of-function mutations (Gerasimavicius et al., 2022). However, some isoform-specific variants will cause other mutational coding effects such as frameshifts, which we have not considered here. Finally, we limited to variants in EICs when mapping disease-variants to alt-exons. We have not considered variants in ENICs - unannotated exons in novel transcripts uncovered by long-read RNA-seq (Shi et al., 2024). Further studies are required to validate and annotate these novel transcripts, and their contribution to proteome diversity.

We anticipate our framework will have broad utility for the interpretation of functional variants for Mendelian diseases and complex traits, to uncover mechanisms of common and rare variants. Assessment of both common and rare disease-associated variants in the context of isoform-specific effects, often through tissue-specific alternative isoforms, will help explain genetic contributions to human disease.

## Methods

### Defining exons classes

Long-read RNA-seq transcripts were obtained from a meta-analysis of long-read RNA-seq from human tissues (Shi et al., 2024). Transcripts associated with coding sequences were retained and exons were classed according to their intersection with the reference or alternative transcripts. The reference transcript was determined from the MANE annotation (Morales et al., 2022), where exons present in the MANE transcript were classed as ref-exon. Exons present in alternative transcripts were classed as alt-exon and further divided into exons that were present in transcripts from Gencode V46, termed exons-in-catalog (EIC). Exons present in alternative transcripts, but not found in Gencode transcripts were termed exons-not-in-catalog (ENIC). The alt-exons were further classed by position in the alternative transcript, into alternative first exons (AFE), alternative last exons, alternative internal exons (including retained intron), and 5’ or 3’ splice site extensions.

### Evolutionary conservation of exons

Conservation was determined by pairwise comparison between human and other species using UCSC Liftover with minMatch 0.5. The fraction of DNA sequences (exons) in synteny was determined by the number of sequences that match as a fraction of total queried sequences. A background model was determined using bedtools shuffle with the following options: -chrom -f 0 -noOverlapping, with repeat elements from UCSC excluded. Five background permutations of bedtools shuffle per DNA sequence set were determined and the mean fraction of syntenic sequences from the five permutations using LiftOver are shown.

Phylop scores were downloaded from the UCSC genome browser. The PhyloP score for exons was determined from the PhyloP470-way imputation. To determine the exon PhyloP score, the mean PhyloP was used from the per nucleotide PhyloP score for each exon using bigWigAverageOverBed from the UCSC genome browser v1.04.00.

### Repeat elements intersection

Repeat elements were obtained from the RepeatMasker track from the UCSC genome browser. Repeat elements were grouped into classes as per UCSC into SINE, LINE, LTR, and DNA repeat elements. Where repeats did not fall into these groups, they were combined and labelled as “other”, which includes low complexity repeats, satellite repeats, and RNA repeats.

To determine the fraction of exons with repeat elements, exon coordinates were intersected with the RepeatMasker track using Bedtools Intersect.

### Exon class variant annotation

Variant occurrence rates were estimated using allele frequencies from gnomAD v4.1, considering only variants that passed quality control in genome and exome sequencing. Variants were binned by allele frequency as follows: common (>0.05 to <0.5), low-frequency (>0.01 to <0.05), and rare (>0.001 to <0.01). For each exon class, the number of variants within each frequency bin was normalised per kilobase to account for differences in total base pair content across classes.

Molecular consequences of variants were annotated using Ensembl VEP version 112 (McLaren et al., 2016), across all coding transcript isoforms available in Ensembl for each exon class. The analysis was limited to exons catalogued by Ensembl. Variant effects were grouped by the “Consequence” category assigned by Ensembl VEP into low (synonymous), moderate (missense), and high impact (stop loss, start loss, stop gain, frameshift). In cases where a variant mapped to multiple transcripts, we considered the most severe consequence.

### SNP enrichment analysis

Background models to estimate enrichment was performed by sampling equal numbers of variants to the condition tested from a union list of variants in the Affymetrix 6.0 and Illumina 650 SNP arrays (n number of total variants). Background SNPs were intersected with features (e.g. alt-exons) and repeated with 10 permutations to obtain a mean background model for each tested condition. Enrichment scores were then computed as the number of observe SNPs (e.g. GWAS category) over the expected background mean over multiple permutations.

### DPP9 peptide analysis

To determine peptide expression from published mass spectroscopy data, DPP9 peptides from the reference isoform (long and short form) were identified using the Global Proteome Machine and Database (GPMDB) (https://www.thegpm.org/GPMDB/index.html) (Fenyö and Beavis. 2015). The top five non-overlapping DPP9 peptides based on the peptide spectrum match of the Westmore-Standing plot for the long and short form DPP9 were used. Human cell line and tissue data for each protein sequence were amalgamated into tissue types.

Tissue Enrichment score for the DPP9-AFE was calculated by using the proportion for each tissue mapped to the DPP9-AFE (observe), over the proportion of tissue associated with the combined tissues from shared (five highest) peptides (expected).

### Phylogenetic analysis

Phylogenetic trees were constructed from BLAST sequences (BLASTn for DNA sequence and BLASTp for protein sequence), allowing up to 5000 hits. DNA sequence identity ranged from 72-100%. Protein sequence identity ranged from 44-100%. Where BLAST sequence matches are find more than once for a given species, the highest percent identity was retained for phylogenetic tree generation. BLAST sequences underwent multiple alignment using ProbCons and alignment processing using Gblocks allowing least stringent alignments. Phylogenetic tree construction was performed by maximum likelihood using PhyML with SH-like approximate Likelihood-Ratio and the substitution model GTR for DNA and Dayhoff for protein. The circular phylogenetic tree was rendered using TreeDyn. Organism silhouette images were taken from PhyloPic v2.0 (link), under Creative Commons licenses.

### Tissue-specific scoring

To determine tissue specificity of GTEx long-read RNA-seq, transcripts per million (TPM) counts were obtained from 22 tissues, with only transcripts with median expression > 0 in any tissue considered. To quantify tissue specificity of mRNA isoforms, the Gini coefficient, a nonparametric statistical measure of inequality, was computed for each transcript, where values close to 1.0 indicate high tissue specificity, while values near 0 reflect constitutive tissues expression. The Gini coefficient was computed as previously described (2, 3), using the ineq package R package v 0.2–13.

### Variant effect prediction in isoforms

To predict variant effects, variants from ClinVar (VCF weekly release 2024-08-05) and the GWAS Catalog were intersected with alt-exon EIC to limit the mapping to alternative isoform-specific exons. Genomic coordinates were first mapped with Ensembl VEP version 112 (29165669, McLaren et al., 2016). Any variant with a non-synonymous consequence in either the ref-exon transcript or the UniProt primary isoform was discarded (UniProt Consortium. 2025). We also discarded variants with a predicted splice consequence in the ref-exon transcript or the primary isoform based on Ensembl VEP annotations, or with a masked SpliceAI score (Jaganathan et al., 2019) (sourced from Illumina BaseSpace Sequence Hub) of >= 0.2. Missense variants were further mapped to alternative transcript isoforms from FLIbase (Shi et al., 2024) with a custom R script, based upon the DNA and protein sequences, the exon boundaries of the transcript, the DNA strand directionality, and the DNA sequence change for the variant. Transcripts predicted to undergo nonsense-mediated decay were excluded. Missense variant effects in the FLIbase-provided protein sequence for the isoforms were predicted with ESM-1v (Meier et al., 2021) as previously described (Livesey and Marsh. 2023). Protein sequences from FLIbase were also modelled with AlphaFold3 (Abramson et al., 2024) using a3m formatted alignments generated with MMseqs2 (Steinegger and Söding. 2017) based on the UniRef and environmental sequence databases. Missense variant effects in the AlphaFold3 models were predicted with FoldX 5.0 (Delgado et al., 2019) by first calling the RepairPDB command on the structure, followed by the BuildModel command to estimate the change in Gibbs free energy of folding (ΔΔG).

### Cell culture

A549 (ATCC, CCL-185) and HEK293T cells were maintained in DMEM (Life Technologies #41965039) supplemented with 10% fetal bovine serum (FBS) and 1% penicillin/streptomycin (P/S). H358 and HCT116 cells were maintained in RPMI or McCoy’s 5A (Gibco # 26600023) media, respectively, each supplemented with 10% FBS, 1% PS. HSAEC1-KT cells were maintained in SABM BulletKit medium (Lonza CC-3119 and CC-4124) containing Bovine Pituitary Extract, hydrocortisone, human Epidermal Growth Factor, epinephrine, transferrin, insulin, retinoic acid, triiodothyronine and fatty acid-free BSA. N/TERT-1 keratinocytes were maintained in keratinocyte-SFM medium (Gibco # 17005042) supplemented with 2.5µg human recombinant epidermal growth factor and 25mg bovine pituitary extract from the same kit. All cell lines were grown at 37°C and 5% CO_2_.

### Constructs and cloning

DPP9-short (Uniprot ID Q86TI2-1) and DPP9-AFE were amplified from H358 cDNA or in-house plasmid constructs using primers Short_HindIII_FW (GCGCAAGCTTATGGCCACCACCGGGACCCCAA) or AFE_HindIII_FW (GCGCAAGCTTATGTTTCATTTCATTTCTCATCT) and DPP9_SalI_RV (GCATGTCGACGAGGTATTCCTGTAGAAAGTGCA) by PCR (98°C for 30s, 30 cycles of 98°C for 10s, 69°C for 20s, 72°C for 130s and 72°C for 3 minutes). Digested and purified products were subcloned into HindIII/SalI-digested pEYFP-N1 (BD Biosciences) to generate in-frame N-terminal YFP-tagged DPP9 isoform expression constructs. The risk variant rs12610495A>G, creating the DPP9-AFE L8>P mutation, was introduced by PCR using primers L8P_FW (CAAAAATTCGATCCCGGGGATGAGAAATGAAATG) and L8P_RV (ATTTCATTTCTCATCCCCGGGATCGAATTTTTG) followed by DpnI template digestion. The catalytically inactive (S729A) mutation was introduced using S729A_FW (CCGTAGGCCCAGCCATGGATGGCAACTCG) and S729A_RV (TGGCTGGGCCTACGGGGGCTTCCTCTCG). The YFP-only control vector was made by excising the DPP9 insert using HindIII and AgeI restriction digestion and re-circularised by T4 ligation with a short linker fragment with compatible sites, made by annealing oligos linker_HindIII_FW (GAAGCTTGGTACCGCGGGCCCGGGATCC) and linker_AgeI_RV (GACCGGTGGATCCCGGGCCCGCGGTACC). All constructs were verified by Plasmidsaurus whole-plasmid sequencing.

### RNA extraction

Total RNA was isolated using the RNEasy Mini Kit from Qiagen (cat.no 74104) according to manufacturer’s instructions including on column DNAse digestion.

### Quantitative RT-PCR

cDNA synthesis was performed using the SuperScriptTM First-Strand Synthesis System for RT-PCR from invitrogen (cat. no. 11904-018) with 1 ug RNA. qPCR was performed in 3 biological replicates with 3 technical replicates each using the LightCycler 480 SYBR Green I Master mix (Roche 04887352001) on the CFX96 Touch Real-Time PCR Detection System (Bio-Rad). The C_t_ values and primer efficiencies (PE) were determined by the CFX Manager software (Bio-Rad) based on a calibration 4-point serial dilution standard curve for each primer pair. The mean relative expression was calculated as PE^−Ct^ and normalized to the geomean of two reference genes, *GAPDH* and *SAFB*. *SAFB* was chosen as it is located on the same chromosome arm as *DPP9* which has heterogeneous copy number in H358 cells. Primers used are listed in supplemental **table S2**.

### Full-Length targEted capture using eXon probes for IsofoRms (FLEXIR-seq)

The full FLEXIR-seq protocol can be found at protocols.io (dx.doi.org/10.17504/protocols.io.n2bvj9dnwlk5/v1) and uses commercially available kits and reagents. Briefly, long, stranded cDNA was prepared from 2ug of total RNA using the ONT cDNA-PCR barcoding kit (SQK-PCS114.24), following manufacturer protocols. RNA underwent reverse transcription and strand-switching with Strand Switching Primer II (SSPII) to generate cDNA. 500ng of cDNA was used for generation of long read libraries using the Twist Mechanical Fragmentation Library Preparation Kit (101280) and the Twist Universal Adapter System (101307). cDNA was subjected to end-repair and dA-tailing using ER/A enzyme, followed by Twist Universal Adaptor ligation (101307), and cleaned using DNA Purification Beads. Amplified indexed libraries were generated by adding Twist Unique Dual Index (UDI) Primers to the adaptor ligated cDNA using KOD Xtreme hot start DNA polymerase (Sigma, (PN 71975-3)), then cleaned with DNA Purification Beads. Library fragments were quantified using DNA Broad range Qubit and fragment size was checked using Fragment analyser. Samples underwent targeted enrichment using Twist Standard Hyb and Wash Kit v2 (104445), Twist Universal Blockers (#100578), and a custom panel of exon probes. Amplified indexed libraries were pooled, then underwent probe hybridisation using the custom exon probe panel, Twist hybridisation mix, and universal blockers, incubated for 16 hours at 70°C. Dynabeads M-270 streptavidin beads (Invitrogen (65305)) in binding buffer were added to the hybridised library, and incubated at 68°C, and immobilised using a magnetic stand, washed, and eluted. Supernatant underwent amplification by PCR and cleaned using DNA Purification Beads. The enriched library was validated and quantified using the Agilent bioanalyser with the High sensitivity assay and a Qubit dsDNA High Sensitivity Assay. The enriched library underwent ONT adaptor ligation to prepare for long-read ONT sequencing using ONT Ligation Kit (V14, SQK-LSK114). Enriched libraries were end-repaired using NEBNext Ultra II end-prep enzyme, followed by adaptor ligation, then AMPure XP bead (Beckman Coulter Agencourt (A63881)) clean-up. Final library was quantified using the Qubit DNA high sensitivity assay. Libraries were sequenced on the PromethION flow cell following manufacturer’s instructions.

FLEXIR-seq was applied to DPP9 isoforms using custom exon probes designed and synthesised by Twist Bioscience, including a probe to the AFE of the DPP9 isoform. To increase library complexity, probes targeting exons of neighbouring genes were included, targeting *MYDGF*, *TNFAIP8L1*, *FEM1A*, *SAFB*, for a total of 327 probes, using the Hg38 genome assembly. Probes have a length of 120 bp, with a mean GC% of 57%.

### Long-read RNA-Sequencing

Target captured samples were multiplexed and sequenced across three flow cells on a PromentION platform (Oxford Nanopore Technologies) using the following software versions: ONT MinKNOW 25.03.7, bream .8.4.4, configuration 6.4.10 and ONT MinKNOW Core 6.4.8. Basecalling was performed using ONT Dorado v.7.8.3 in high-accuracy mode.

### FLEXIR-seq Read processing, demultiplexing and filtering

Reads passing the minimum Q9 score were demultiplexed based on i5 and i7 dual indexes using a custom python script. Adapters were trimmed using cutadapt v.4.6 (Martin et al 2011) with the parameters --report=minimal -m 100 --rc -e 0.12 -j 0 to search for adapters in both forward and reverse orientations, allowing a maximum error rate of 0.12 while filtering out reads shorter than 100bp. Read quality was assessed prior to trimming using NanoPlot v.1.44.0 from the NanoPack2 package (De Coster and Rademakers, 2023). Nucleotide diversity at the read start and end were assessed NanoQC v.0.10.0 (De Coster et al 2018) both prior to and following adapter trimming to confirm successful trimming. Read quality was compared after and before adapter trimming using NanoComp v.1.24.0 (De Coster et al 2018).

### FLEXIR-seq Alignment

Demultiplexed, trimmed reads were aligned to the GRCh38 human genome assembly using minimap2 v.2.28-r1209 (Li. 2018) with the parameter -ax splice for splice junction aware mapping of cDNA reads, achieving >99.6% mapping across all samples. Sam files were converted to bam format, sorted and indexed using SAMtools v.1.20 (Danecek et al 2021). Mapping statistics were generated using SAMtools flagstat command. Aligned bam files for each sample were visually inspected over the DPP9 region in the IGV genome browser (Thorvaldsdóttir, Robinson and Medirov 2013).

### FLEXIR-seq Gene and transcript quantification

Gene and transcript isoform counts were generated using IsoQuant v.3.7 with the parameters -d ont --report_novel_unspliced true --sqanti_output --genedb to enable de-novo isoform discovery optimised for Nanopore sequencing reads. The GRCh38 human reference genome assembly was provided as --reference and the GRCh38 human gene annotation in gtf format, additionally containing all DPP9 transcripts annotated in FLIbase (Shi et al 2024), provided as --genedb. Comparisons with short-read RNA-seq data used the following GEO Sequence Read Archive DataSets: A549 (SRR32141251, SRR32141252, SRR32141253); H358 (SRR24149739, SRR24149740, SRR24149741); HBEC3 (SRR32905582, SRR32905583, SRR32905584); HSAEC1-KT (SRR28773308, SRR28773309, SRR287733010); N/TERT-1 (SRR19142533, SRR19142536, SRR19142539). Short-read RNA-seq reads were aligned using bowtie2 v2.5.3 to GRCh38, and mapped reads converted to gene counts using featureCounts v2.1.1, then normalised using transcripts per million (TPM).

### DPP9 isoform cell transfection and protein extraction

A549 cells at 70% confluence (1.5×10^6^cells) in 6-well plates were transfected with DPP9 constructs (Abbott et al., 2000, Ajami et al., 2004) using Lipofectamine™ 3000 (Invitrogen, Carlsbad, CA; L3000015) following manufacturer’s instruction. 24 hours post-transfection, cells were washed twice with PBS (Gibco, Paisley; #10010023) and then harvested using TrypLE (Gibco, #12605028). Briefly, cells were treated with 0.5 mL of TrypLE for 5 minutes (mins) at 37°C, neutralised in 1 mL of complete media, and spun down at 300 g for 7 mins at room temperature, then the pellet was washed twice with ice-cold PBS.

For whole cell lysate, cell pellets were resuspended in Lysis Buffer (50 mM Tris-HCl pH 7.6, 1% Triton X-100, 10% glycerol, 2 mM dithiothreitol (DTT), 1 mM EDTA) and 1X protease inhibitor cocktail (Roche, Mannheim, #11836170001) for 5 mins on ice and then spun down at 10,000 g for 5 mins. Supernatant containing soluble protein lysate was stored at −30°C.

For nuclear and cytosolic fractions, cell pellets were first resuspended in swelling buffer (10 mM HEPES pH 7.6, 10 mM KCl, 1 mM DTT, 1 mM EDTA, 0.1% Triton X-100 with protease inhibitors) and incubated on ice for 5 mins to swell up. Then, the solution was resuspended several times to disrupt the cell membrane. Samples were spun down at 2000 g for 5 mins at 4°C and supernatant containing the cytosolic fraction was collected. Glycerol was added to a final concentration of 10%. The nuclear pellet was washed in swelling buffer (excluding Triton X-100) and then resuspended in 20 mM Tris-Cl, 420 mM NaCl, 1.5 mM MgCl_2_, 0.2 mM EDTA, 25% glycerol and protease inhibitors. Salt concentration was adjusted using 5 M NaCl to disrupt the nuclear membrane. Nuclear lysate was vortexed and then spun down at 10,000 g for 10 mins at 4°C. Supernatant containing soluble nuclear proteins was collected and stored at −30°C.

### DPP9 enzymatic assay

Protein lysates from A549 cells (1.5×10^6^) were diluted 1:100 in triple distilled water (TDW) and then quantified using Pierce MicroBCA assay (Thermo Scientific (23235)) following the manufacturer’s instructions. To measure DPP9 enzyme activity, two assays were employed. The substrate H-Gly-Pro-ρ-nitroanilide (H-Gly-Pro-pNA, Bachem (4025614)) is hydrolysed by DPP4, 8 and 9 into pNA. One assay used sitagliptin, a potent selective DPP4 inhibitor, to measure DPP8/9 activity, as described previously (Yu et al., 2009). The second assay used sitagliptin and compound 42 (Cpd42), a novel selective DPP9 inhibitor (Benramdane et al., 2023), to measure DPP9 activity by subtraction. Each assay used 15 µg of protein of cell lysate and was performed at pH 7.8 and 37°C. DPP9 S729A/S729A gene knock-in mouse embryonic fibroblast (GKI MEF) cells generated previously (Gall et al., 2013) were a negative control.

To exclude DPP4 activity, protein samples were incubated in 1 mM DTT, 1 µM sitagliptin in a total of 50 µL Tris-EDTA (TE, Sigma-Aldrich (T9285)) buffer on ice for 10 mins. Absorbance at 405 nm (for pNA) and 570 nm (background) were then measured as time zero then 50 µL of 2 mM H-Gly-Pro-pNA was added. Absorbances were measured every 5 mins for 1 hour (Yu et al., 2009). For the second assay, 1 µM of Cpd42 was used in addition.

To calculate enzyme activity, background absorbance at 570 nm was subtracted from absorbance at 405 nm. pNA product was extrapolated, using a pNA standard curve, as µmol pNA, and lysis buffer control values were subtracted. To calculate DPP9 enzyme activity, pNA product from the 2nd assay was subtracted from the DPP8/9 measurement from the first assay. Enzyme activity was then converted to picomole pNA per min per µg of protein lysate.

Enzyme kinetics assays were performed using the following dipeptide substrates coupled to the fluorogenic cleaving group 7-amino-4-methylcoumarin (AMC): H-Valine-Proline (Enzo Life Sciences (BML-P448)), H-Tryptophan-Proline (Enzo (BML-P450)), H-Lysine-Proline (MedChemExpress (P4426A), and H-Aspartate-Proline (Mimotopes, customised).

Substrates were dissolved in 100% DMSO as 2 mM stock solutions. A titration of cell lysate determined the quantity (2.5 µg protein) to use in the kinetics assay, in a range of substrate concentrations. To exclude DPP4 activity, 1 µM sitagliptin was added. Each assay included at least three technical replicates and three biological replicates. Values from blank controls were subtracted from the final readings. The rate of hydrolysis, Δfluorescence/min, at each substrate concentration was calculated and plotted in GraphPad Prism V9.0. Vmax, K_half_ and Hill coefficient were calculated using allosteric sigmoidal equation.

### Immunoblotting

Immunoblot methodology was adapted from (Yao et al., 2011). Approximately 15 µg of protein lysate was mixed with 1X NuPAGE LDS Sample Buffer (Invitrogen (NP0007)) and 1X NuPAGE Sample Reducing Agent (Invitrogen (NP0009)) and TDW to a total volume of 20 µL. Samples were incubated at 95°C for 5 mins and then cooled on ice. PageRuler Plus Prestained Protein Ladder (Invitrogen (26619)) and samples were loaded onto NuPAGE 4-12% Bis-Tris, 1.5 mm Mini Protein gel (Invitrogen (NP0336)) and run with 1X NuPAGE MOPS Running Buffer (Invitrogen (NP001)) at 150V for 1 hour at room temperature.

Proteins gels were transferred onto PDVF membrane by wet transfer, using NuPAGE 1X Transfer Buffer (Invitrogen (NP0006)) at 20V, 300 mA for 1 hour at room temperature. Blots were blocked in 5% skim milk and PBS-T (PBS, 0.5% Tween-20) for 1 hour at room temperature then incubated with anti-DPP9 antibody (Abcam (ab42080)) overnight at 4°C and then goat anti-rabbit horseradish peroxidase (HRP) (Agilent (P044801)) for 1.5 hours at room temperature. Blots were imaged using Immobilon Chemiluminescent HRP substrate (Merck (WBKLS0500)) and a BioRad Chemidoc Imaging System. HRP-conjugated antibody to beta-actin (Abcam, ab49900) was used for the loading control.

Protein expression was analysed using Fiji software (Schindelin et al., 2012). Briefly, mean grey value of each lane was calculated by inverting the image and then subtracting the background. Then, protein expression was normalised by dividing the DPP9 bands by the β-actin band.

## Data availability

FLEXIR-Seq data has been deposited in the Genome expression Omnibus (GEO) respository under the accession number GSE303335. Variant data with exon class files, variant effects, and AF3 predicted isoform protein models will be made available upon publication.

## Code availability

No codes have been generated in this manuscript. Data analysis used existing code.

## Author contributions

G.W.., M.B.., E.T.F.., J.M.H.N., contributed to data curation, visualization and analysis. A.I., B.J.L., A.C., E.F.H., M.W., and J.S. contributed to data analysis and computational resources. J.K.B., C.S. and V.V. contributed data curation. G.W. A.C., S.C.B., and L.M. developed the FLEXIR-seq methodology, and data analysis. J.A.M., W.A.B., and S.C.B. contributed to conceptualization. M.D.G., J.A.M., W.A.B., and S.C.B. contributed project supervision. E.T.F., and M.D.G. reviewed and editing the manuscript. G.W., M.B., J.M.H.N., J.A.M., W.A.B., and S.C.B. wrote the manuscript.

## Supplemental Figure Legends

**Figure S1.**
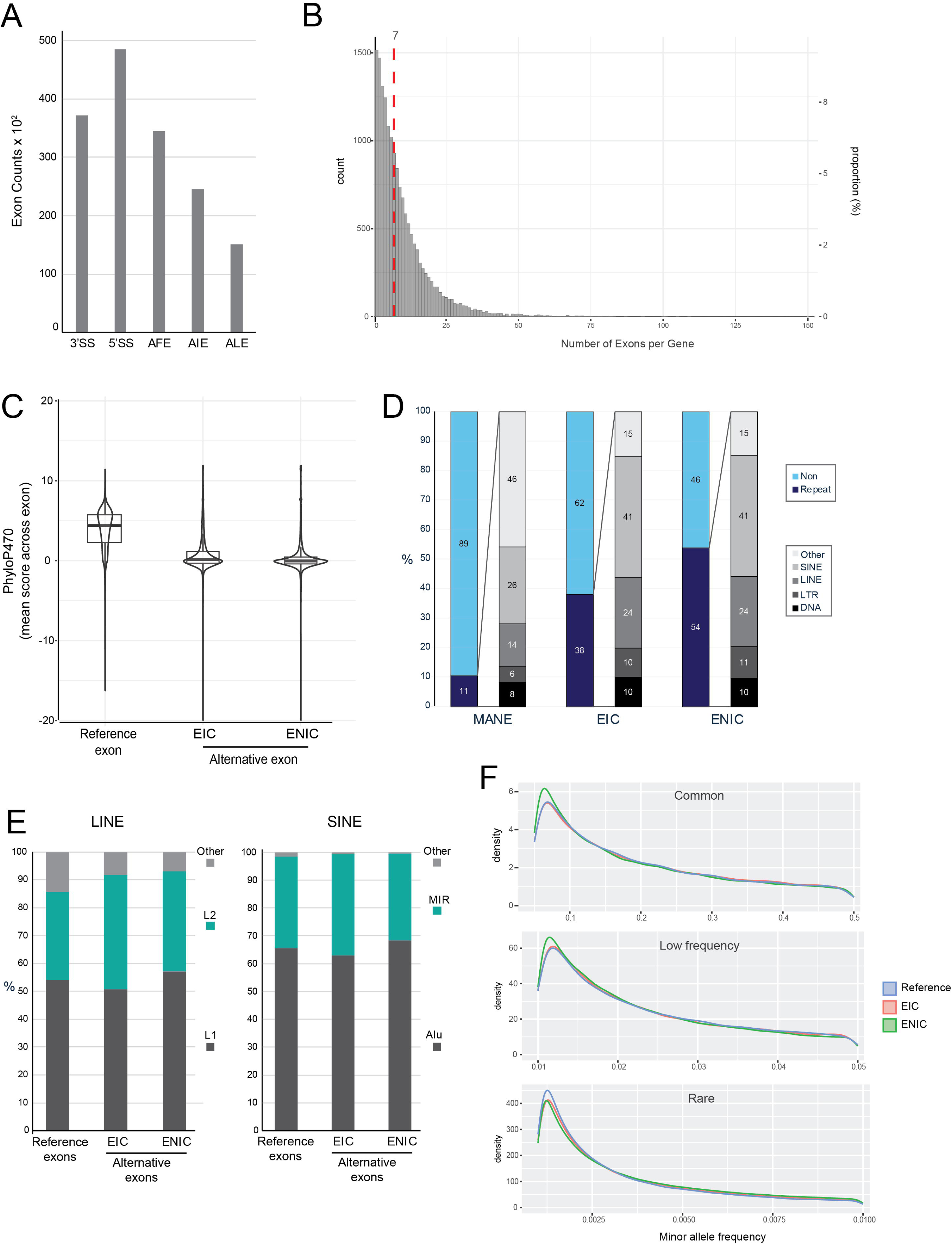
Characteristics of ref-exon and alt-exon classes. **A.** Bar graph of alt-exons in catalogue (EIC) by alternative exon type: 3’ splice site (3’SS), 5’ splice site (5’SS), alternative first exon (AFE), alternative internal exon (including retained introns), and alternative last exon (ALE). **B.** Histogram of exon counts per gene, using all exons in any transcript isoform. **C.** Violin and boxplot of conservation score of exon classes using the mean 470-way PhyloP score across each exon. **D.** Proportional bar chart of repeat element types for the exons per class that intersect repetitive sequences. **E.** Proportional bar chart for sub-types of LINE and SINE transposable repetitive elements per exon class. Only the two most common sub-elements are shown, with remaining sub-elements grouped as ‘other’. **F.** Density distribution plots of single nucleotide variant (SNV) counts from gnomAD by exon class, across the allelic frequencies for minor allele frequency groups.

**Figure S2.**
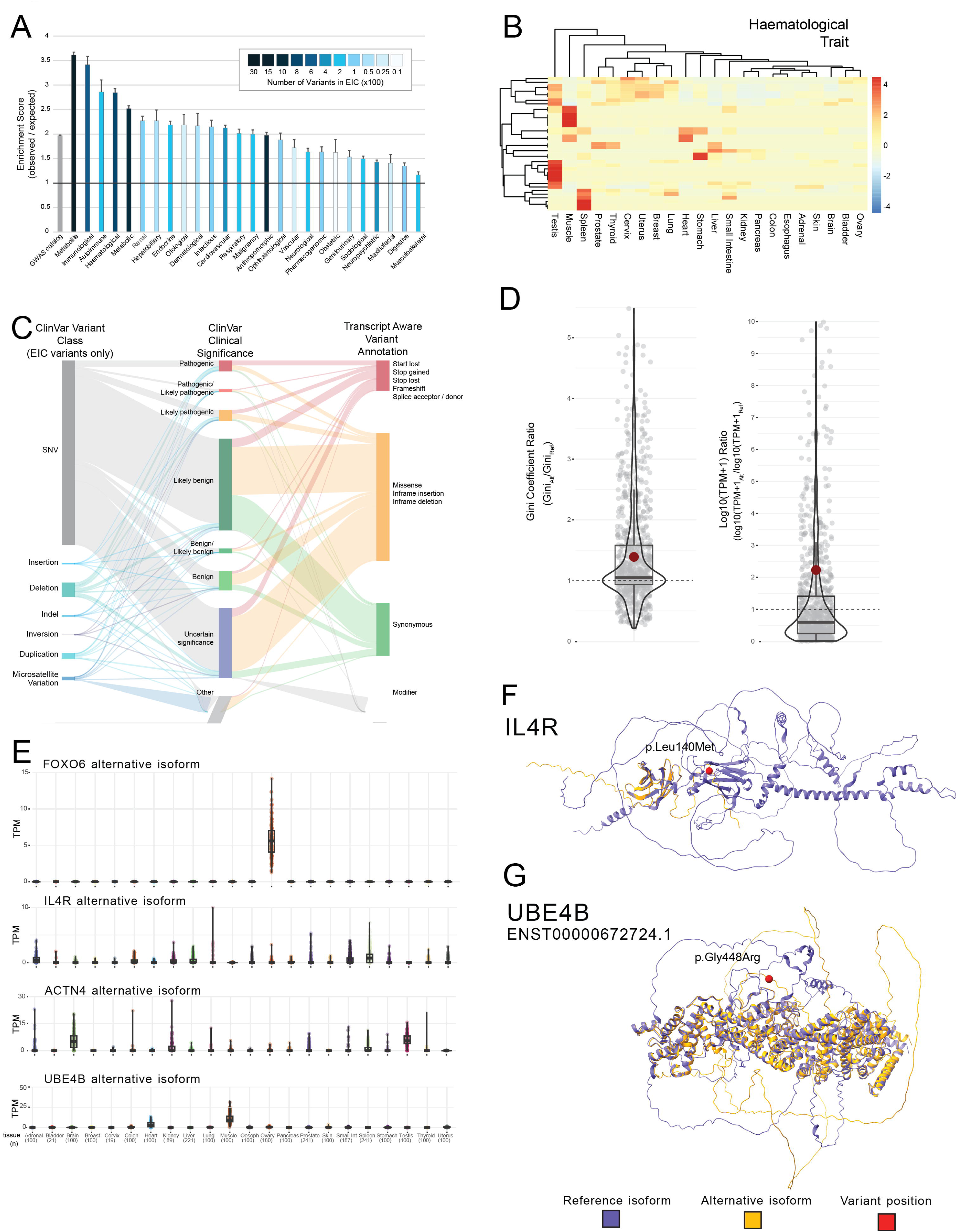
Common and rare variant association with alternative transcript isoforms. **A.** Enrichment score for GWAS catalog variants in non-canonical exons in catalogue. Grouped GWAS catalog variants associated with non-canonical exons in catalogue (observed), with scores computed from ten imputations of randomly selected genomic variants (expected), using equivalent numbers of trait group variants. **B.** Heatmap of alternative transcript isoform expression for isoforms associated with variants from haematological GWAS traits. Expression is from long-read RNA-seq transcripts in 22 GTEx tissues. Expression as z-score of log10 TPM. Hierarchical clustering for tissues and variants was performed using the ward method. **C.** Proportional Sankey plot of ClinVar variants associated with non-canonical exons in catalogue by variant class, ClinVar annotated significance, and transcript isoform-aware variant annotation. **D.** Box and violin plots of alternative isoform to reference isoform ratios for tissue-specificity metric (Gini coefficient ratios, left), and transformed expression (log10(transcripts per million (TPM)+1) ratios, right). Benign, likely benign, and variants of uncertain significance (VUS) ClinVar variant-mapping alternative isoforms and gene-matched reference isoforms were annotated with Gini coefficients and TPM values from long-read RNA-seq of 22 GTEx tissues. For transformed TPM, ratios were determined from the highest expression in any GTEx tissue. Where a ClinVar variant maps to multiple transcripts, only the alternative transcript isoform with the most damaging VEP score was used. Mean is shown in red. **E.** Reference isoform (blue) and variant-associated alternative isoform (yellow) for IL4R. GWAS variant position shown in red. **F.** Reference isoform (blue) and variant-associated alternative isoform (yellow) for UBE4B. GWAS variant position shown in red. **G.** Violin and boxplot for alternative transcript isoform expression across 22 GTEx tissues from long-read RNA-seq.

**Figure S3.**
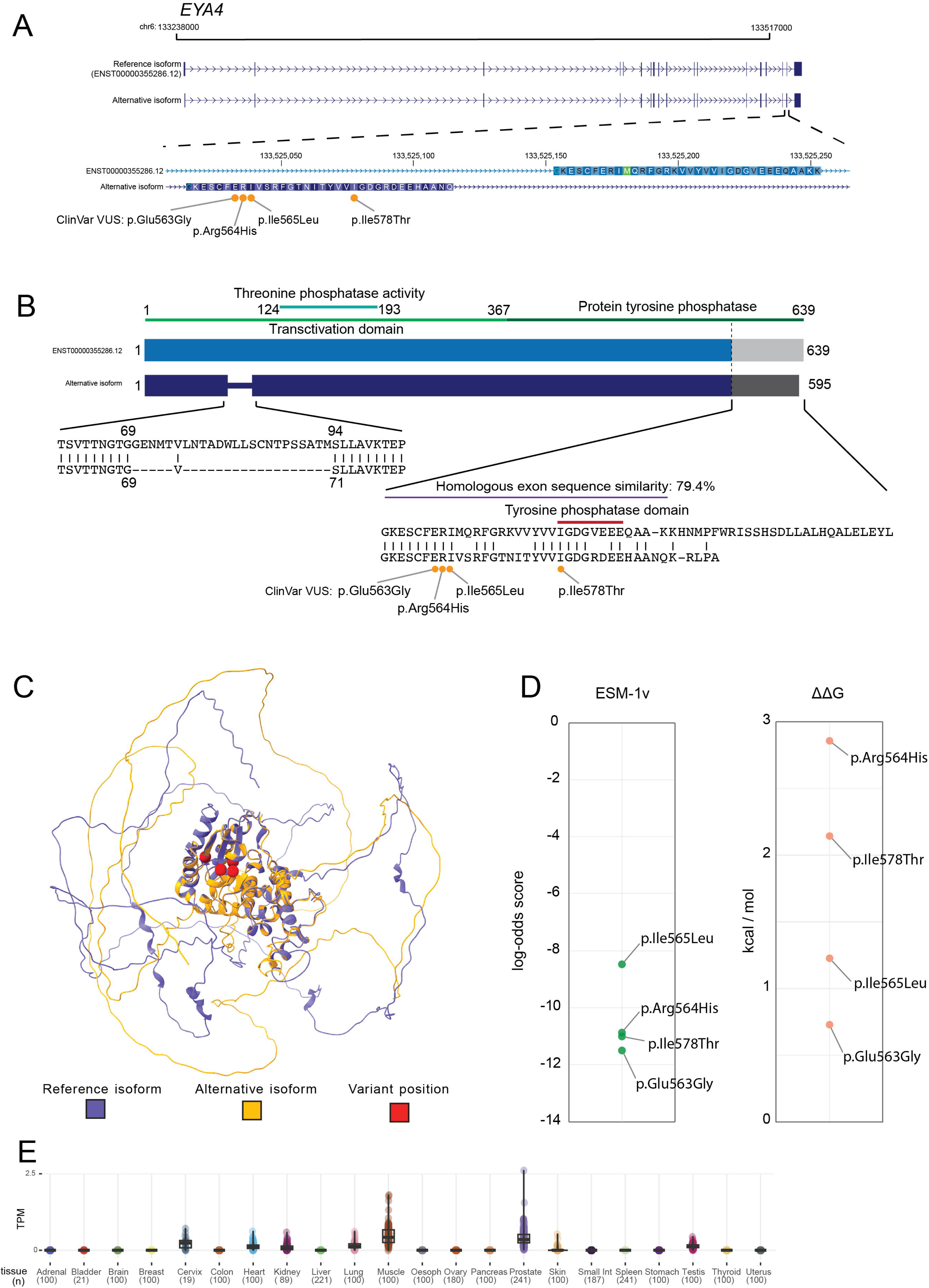
Variants of uncertain significance in an alternative homologous exon of an EYA4 isoform. **A.** *EYA* transcript isoforms, showing the reference isoform (ENST00000355286.12), and an unannotated alternative isoform. The 3’ end of the *EYA4* gene has two adjacent homologous exons that show mutually exclusive splicing. Four ClinVar missense variants of uncertain significance (VUS) map to the alt-exon in the alternative transcript isoform. **B.** The EYA4 isoform protein contains multiple functional domains, with shared isoform protein sequences, including the peptide sequence encoded by the homologous exons. **C.** Predicted AF3 structure for the reference (purple) and alternative (yellow) EYA4 isoforms, with the ClinVar VUS indicated in red. **D.** Variant effect prediction for four missense VUS in the alternative EYA4 isoform using evolutionary scale modelling (ESM)-1v, and FoldX-derived thermodynamic stability difference in the Gibbs free energy (ΔΔG) ranked scores. **E.** Violin and boxplot for *EYA4* alternative transcript isoform expression across 22 GTEx tissues from long-read RNA-seq.

**Figure S4.**
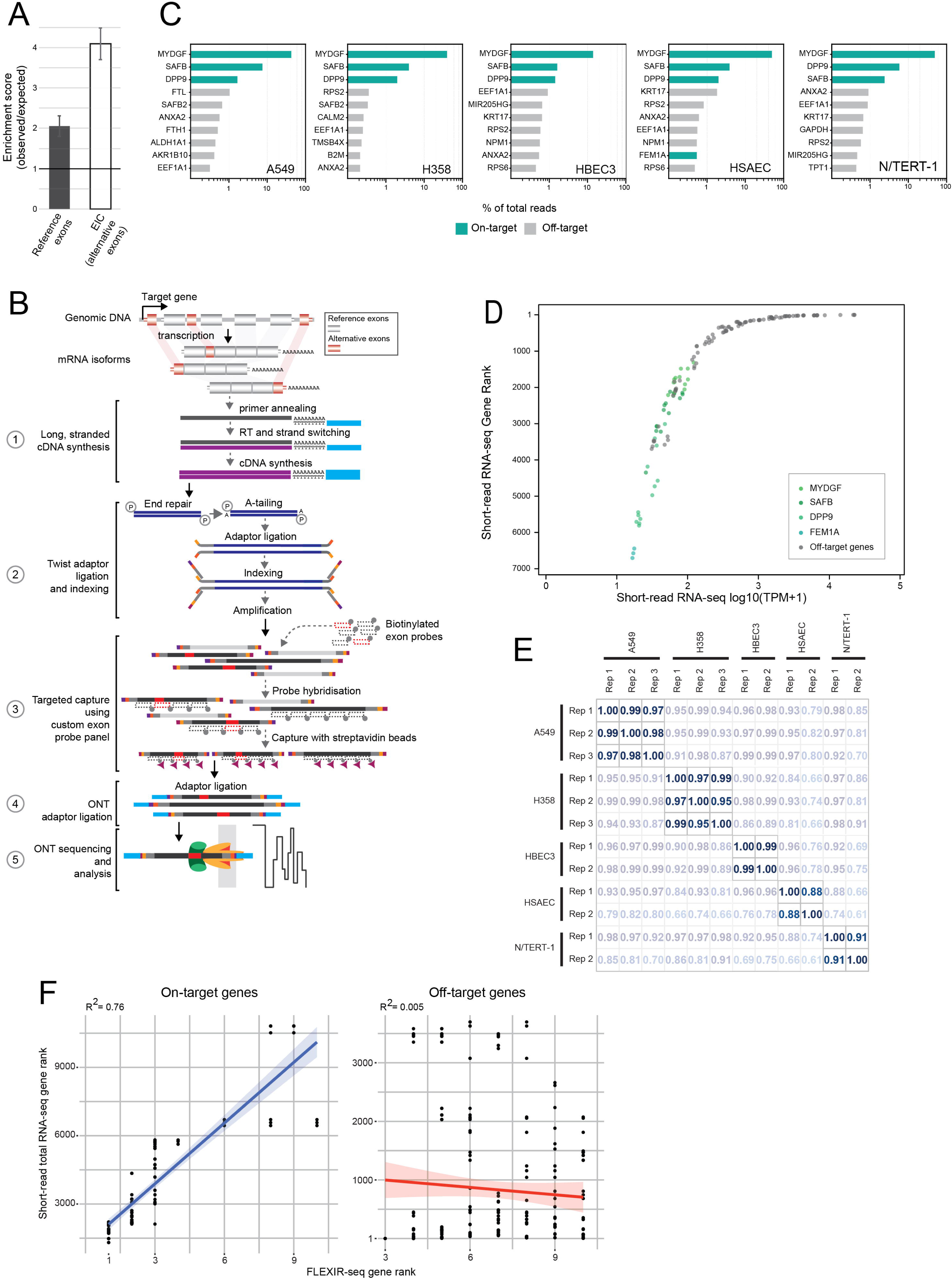
Full-Length targEted capture using eXon probes for IsofoRms (FLEXIR-seq) **A.** Enrichment of Severe COVID-19 lead variants and 95% credible set variants in ref-exons and alt-exons in catalogue (EIC). Equal number of randomly selected genome variants to the observed variant in exon classes were used as the expected occurrence, with ten imputations for 95% confidence intervals of the mean score. **B.** Schematic for FLEXIR-seq protocol, using a custom panel of biotinylated exon probes to capture full-length cDNA for long-read RNA sequencing. **C.** Top 10 ranking genes from FLEXIR-seq gene panel in five cell lines with pooled replicates, with on- and off-target genes based on targeted exon probe design. Transcript isoforms for each gene were combined per gene and calculated as percentage of total sequenced reads. **D.** Scatter plot of short-read total RNA-seq gene ranks against transcripts per million (TPM, normalised as log_10_(TPM+1) for on-target and off-target genes identified using top 10 genes from FLEXIR-seq. Short-read total RNA-seq data were from three replicates for each cell line. **E.** Pearson coefficients for biological replicates of cell lines, using correlations of full-length transcripts for *DPP9* expressed as percentage of total reads for each replicate. **F.** Linear model fit of the top 10 genes from FLEXIR-seq using gene ranks from replicates against gene ranks from short-read total RNA-seq for three replicates for each cell line. (Kavaliauskas et al., 2024). Linear model fits were generated separately for on-target genes (left and off-target genes (right).

**Figure S5.**
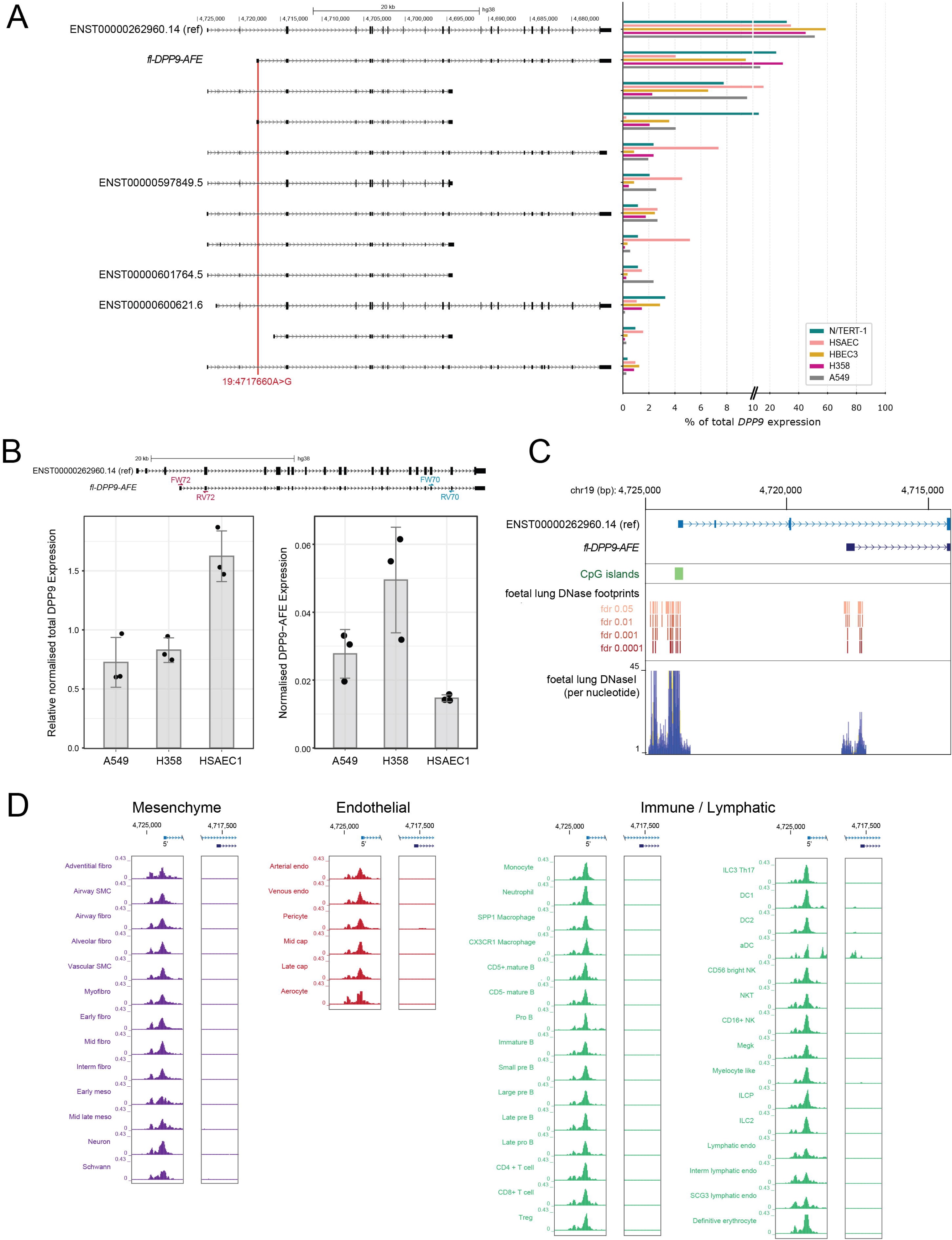
A DPP9 transcript isoform is associated with a severe COVID-19 variant. **A.** *DPP9* transcript isoforms from targeted long-read RNA-seq (FLEXIR-seq) in five cell lines. Transcript expression is shown as percentage of total *DPP9* expression. **B.** Schematic showing primer annealing sites for forward (FW70) and reverse (RV70) primers used to amplify all *DPP9* transcripts (left panel) and for forward (FW72) and reverse (RV72) primers used to amplify the DPP9-AFE isoform (right panel). For all *DPP9* mRNA isoforms (left) or the *DPP9-AFE* mRNA isoform (right), expression was normalized to *GAPDH* and *SAFB* expression in A549, H358 and HSAEC1 lung epithelial cell lines. **C.** Genome browser image of the reference transcript promoter, and *fl-DPP9-AFE* alternative promoter, showing location of CpG islands, and ENCODE DNaseI hypersensitive sites and footprints (at various false discovery rates; FDR), from foetal lung tissue (Vierstra et al., 2020) **D.** Genome browser image of published single-cell ATAC-seq tracked for mesenchyme, endothelial and immune/lymphatic cell types from foetal human lung. The pseudo-bulk ATAC-seq signal for the promoter of the reference transcript (left) and alternative promoter of the 19:4717660A>G variant-associated transcript isoform (right) are shown.

**Figure S6.**
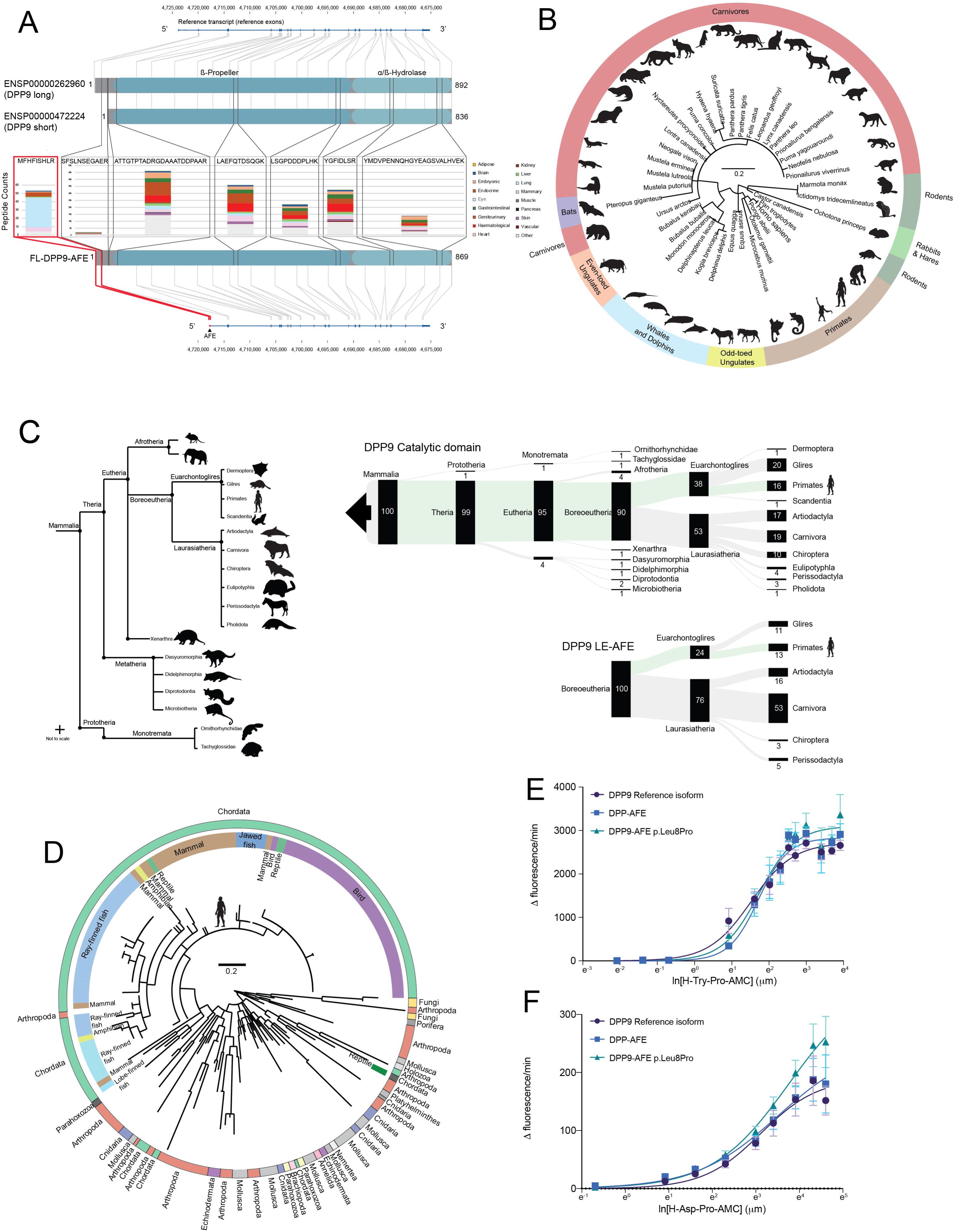
DPP9 protein expression and evolutionary conservation. **A.** DPP9 peptide expression, including the AFE, and five highest detected peptides from proteomic meta-analysis (Fenyö and Beavis. 2015) used to generate the background model. **B.** Phylogenetic tree from maximum likelihood analysis of DPP9 alternative first exon peptide sequence matches from BLASTp. **C.** Phylogenetic tree of mammalian clades associated with the DPP9 peptide phylogenetic ranges, and phylogenetic range of the DPP9 catalytic domain, with DPP9 alternate isoform alternative first exon encoding peptide. Clades determined from BLASTp using maximum likelihood analysis. The catalytic domain clades extend beyond Mammalia (figure S6D). Green indicates the clades to *Homo sapiens*. Numbers represent percentages. **D.** Phylogenetic tree from maximum ikelihood analysis of DPP9 catalytic domain peptide sequence matches from BLASTp. **E** and **F**. DPP9-specific enzyme kinetics for DPP9 reference and AFE isoforms, and the p.Leu8Pro variant form of DPP9-AFE, using H-Try-Pro (E) and H-Asp-Proline (F) peptides. Fluorescence emission was detected upon cleavage of the fluorogenic salt 7-amino-4-methylcoumarin (AMC), then the rate of hydrolysis data fit to an allosteric sigmoidal model.

